# Intestinal fructose metabolism drives unsaturated fat absorption and synergizes with GLP-1 receptor agonism to promote weight loss

**DOI:** 10.64898/2026.06.03.729910

**Authors:** Carolina E. Echeverría, Mujmmail Ahmed, Jonathan Gao, Sarah L. Stewart, Isaac Nathoo, Lucas K. Debarba, Carlos A. Lafourcade, Tanvir Ahmed, Odinakhon Shamieva, Tiffany Perrier, Chandana Prakashmurthy, Anthony Escamilla, Philip Moon, Jeshua Kim, Rachel K. Zwick, Lewis C. Cantley, David E. Cohen, Marcus D. Goncalves

## Abstract

High-fat, high-sucrose (HFHS) diets are established risk factors for obesity. In the intestine, sucrose is hydrolyzed into glucose and fructose, with fructose being taken up by epithelial cells and phosphorylated by ketohexokinase (KHK). We hypothesized that KHK is required for the obesogenic effects of HFHS diets and performed genetic and pharmacologic experiments in mice using diet-induced obesity (DIO) models. We show that genetic loss of KHK prevents HFHS-induced weight gain and intestinal villus elongation. Moreover, pharmacologic inhibition of KHK (KHKi) promotes weight and fat loss during continued HFHS feeding in DIO mice and enhances weight loss and weight maintenance during and after incretin-mimetic therapy. The anti-obesogenic effects of KHKi were associated with delayed intestinal lipid absorption, reprogramming of lipid metabolism in the distal intestinal epithelium, and reduced absorption of unsaturated dietary fats. Together, these findings identify fructose metabolism as a key regulator of intestinal lipid handling and suggest that fructose promotes obesity, in part, by enhancing intestinal lipid absorption and metabolism.

## Introduction

Western style diets are characterized by the combined intake of high amounts of refined sugars and dietary fat, a macronutrient pairing that promotes weight gain more potently than either component alone^1,2^. The widespread incorporation of both sugar and fat into processed foods has paralleled the rise in obesity and metabolic dysfunction among Americans over the past five decades^3^. Notably, all commonly consumed caloric sweeteners contain significant proportions of fructose, including table sugar (sucrose) (50% fructose), honey (40% fructose), and high-fructose corn syrup (HFCS) (55% fructose)^4^. Although fructose has long been present in human diets, contemporary intake levels in Western populations are now more than ever recorded^5^. Beyond its caloric contribution, fructose exhibits distinct metabolic effects compared with glucose, including preferential first pass metabolism, alterations in lipid handling, and promotion of systemic lipid storage under conditions of combined sugar and fat intake^6,7^.

Within the body, dietary fats and sugars are absorbed via the intestinal villi, fingerlike epithelial projections that line the small intestine and increase nutrient absorption. The small intestine is anatomically and functionally divided into at least three regions: duodenum, jejunum, and ileum^8,9^. Dietary fats and complex carbohydrates are initially digested in the duodenum, which contains high concentrations of pancreatic enzymes and bile acids required for lipid emulsification^10^. However, the majority of dietary lipid and monosaccharide absorption occurs in the jejunum, a region characterized by high transporter expression, rapid nutrient uptake kinetics, large enterocyte cytoplasmatic lipid droplets (CLDs), and robust lymphatic lipid output, as compared with the ileum^11^.

At physiologic intake levels, the small intestine serves as the dominant site of first-pass fructose clearance, converting fructose into glucose and organic acids and limiting hepatic exposure until intestinal capacity is saturated^12,13^. Genetic and tracer-based studies indicate that intestinal carbohydrate sensing and fructose metabolism are actively regulated processes that determine systemic fructose delivery and tolerance^13,14^. The specific effects of fructose on the intestinal epithelium structure, lipid handling, and absorptive efficiency in the context of obesity remain incompletely understood.

In the enterocyte, fructose uptake via Glut-5 is phosphorylated by Ketohexokinase (KHK) to form fructose 1-phosphate (F1P) where it can be metabolized further to fuel glycolysis and de novo lipogenesis^15,16^. We have previously demonstrated that mice exposed to HFCS develop longer intestinal villi and improved nutrient absorption, as compared to control mice^2^. Others have shown that intestinal KHK-C overexpression enhances first-pass fructose clearance and shields the liver from fructose-induced lipogenesis, whereas deleting intestinal KHK-C allows fructose to bypass the gut and reach the liver intact^7,15^. Whole-body KHK deletion protects mice from fructose-induced obesity, but this protection has been attributed primarily to reduced hepatic lipogenesis and altered food intake^7,17,18^. Whether fructose metabolism within enterocytes directly regulates intestinal lipid handling, independent of villus elongation or hepatic effects, has not been explored.

Over the last decade, the development of pharmacologic therapies for weight loss has expanded rapidly, with major advances in the treatment of obesity and type 2 diabetes^19^. Potent glucagon-like peptide-1 (GLP-1) receptor agonists (GLP1RAs), such as semaglutide, safely and effectively reduce body weight and improve obesity-related complications^20,21^. Combination approaches that pair GLP1RAs with complementary therapies can further enhance fat mass loss^22^. However, sustained treatment is required to maintain these benefits and prevent weight regain^23^, highlighting the need for alternative strategies that improve long-term tolerability, durability, or transition to maintenance therapies^24^. To date, no studies have evaluated the combination of pharmacologic KHK inhibition (KHKi) with GLP1RAs.

Here, we show that mice on a HFHS diet rapidly accumulate fat mass and that this effect is blocked by genetic KHK deletion or reversed by KHKi in established obesity. KHKi also enhances semaglutide-induced fat loss and delays weight regain following GLP-1RA withdrawal. Mechanistically, we find that fructose metabolism reprograms enterocyte lipid handling, promoting CLD formation and proximal intestinal fat absorption, independently of changes in villus architecture. These findings identify intestinal fructose metabolism as a previously unrecognized regulator of dietary lipid absorption and suggest that combining KHK inhibition with incretin-based therapies may improve outcomes in obesity management.

## Results

### High-fat high-sucrose diet increases fat accumulation in a ketohexokinase dependent manner

To study the impact of diets high in fat and sugars in the context of weight gain and rapid fat accumulation, we generated a series of customized diets that contain either sucrose or dextrose (D-glucose) as the primary source of sugar. These diets were designed to be representative of the typical American diet with 45%, 30% and 25% of energy from fat, carbohydrates, and protein, respectively^25^. Healthy male and female C57BL/6J (KHK^+/+^) mice were fed either a low-fat (LF), a high-fat high-sucrose (HFHS) or a high-fat high-dextrose (HFHD) diet. Over a period of 5 weeks, mice on the HFHS diet gained more body weight and body fat than mice eating the LF diet without changes in lean mass (**Figures 1A, 1B, and S1A**). The HFHD diet also increased weight and body fat but to a lesser extent than HFHS. The accumulation in body fat for both high fat diets was detectable as early as seven days after diet initiation and continued to rise through the five-week period. We noted a transient increase in food intake the first few days after HFHS diet initiation, but this effect normalized over time despite continued weight gain (**Figure S1B**), consistent with prior reports^18,26^. The excess body weight and fat mass observed in the HFHS group correlated with increases in the masses of the perigonadal white adipose tissue (gWAT), but not gastrocnemius (**Figure S1E, F**). Similarly, the HFHD group showed increased gWAT mass, although to a lesser extent than sucrose. Similar results were observed in female mice (**Figure S2**). Together, these findings demonstrate that high fat high sugar diets promote rapid weight and fat accumulation in mice, with sucrose exerting a more pronounced effect than dextrose, despite equivalent energy intake.

**Figure 1.**
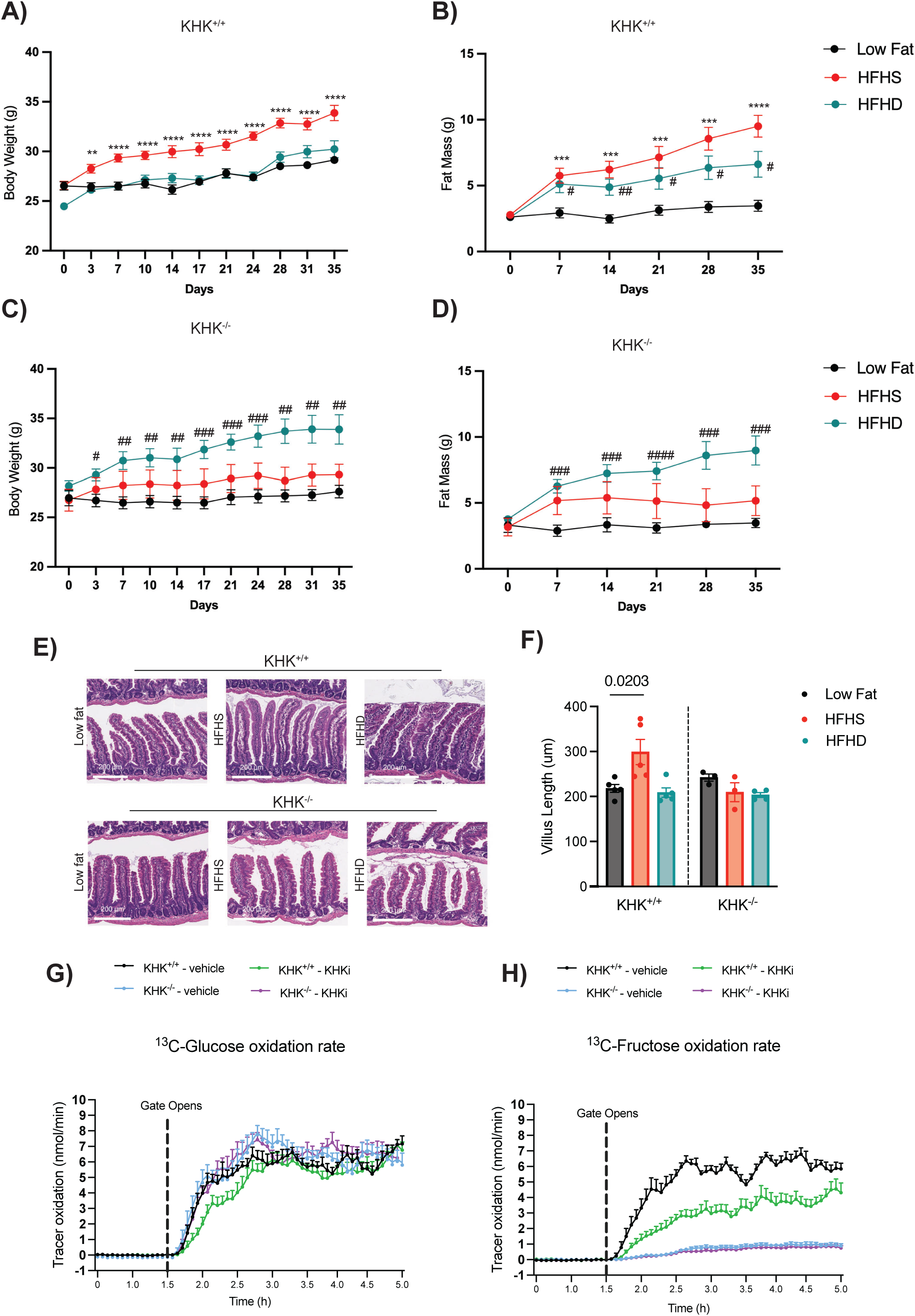
High-Fat High-Sucrose diet leads to a fast fat accumulation over time: **A)** Body weight over time of C57BL/6J male KHK^+/+^ were fed either low fat diet (Black), high-fat high-sucrose (HFHS; red) or high-fat high-dextrose (HFHD; teal) diets for five weeks. **B)** Fat mass over time in male KHK^+/+^ mice measured by echoMRI (n=10 mice per group). **C)** Body weight over time and **D)** Fat mass over time in male KHK^-/-^ mice under the same experimental conditions described above (n= 9 mice per group). **E)** Representative hematoxylin and eosin (H&E)-stained duodenum from either KHK^+/+^ or KHK^-/-^ after five weeks on low Fat, HFHS or HFHD diets; Scale bar: 200 µm. **F)** Quantification of duodenal villus length from histological sections (n=3-5 mice per group). Time-course of whole-body **G)** ^13^C-glucose and **F)** ^13^C-fructose oxidation rates measured over 5 hours in KHK^+^/^+^ and KHK^-^/^-^ mice treated with vehicle or KHKi (ketohexokinase inhibitor) under a HFHS diet (n= 8 mice per group). Tracer oxidation is expressed as nmol/min. Data are presented as mean ± SEM with individual data points overlaid. *P* values were calculated using two-way analysis of variance (ANOVA) with Tukey’s multiple comparison test. *^/#^ *p*<0.05 **^/##^ *p*<0.01, ***^/###^ *p*<0.001, ****^/####^ *p*<0.0001.*(Low Fat vs HFHS), ^#^(Low Fat vs HFHD).

KHK-dependent fructose phosphorylation has been shown to be necessary for sugar-induced metabolic disease^17,27,28^. To test the role of KHK in the response to the dextrose- and sucrose-rich high fat diets, we fed the same dietary interventions described above to male and female mice with whole body genetic KHK knockout (KHK^-/-^)^29^. KHK^-/-^ mice were largely protected from HFHS-induced weight gain, fat accumulation, and hyperlipidemia (**Figures 1C, 1D, S2, Table 1**). Conversely, the HFHD diet increased body weight, fat mass, and gWAT mass, as compared to the LF or HFHS counterparts (**Figures 1C, 1D, and S1E**). These effects correlated with increased food intake in KHK^-/-^ mice in both HFHS and HFHD, as compared to their LF group (**Figure S1D**). While there were no changes in lean mass over time in any diet group, the HFHD cohort started with more lean mass despite random group assignment (**Figure S1D**).

**Table 1:** Effects of HFHS feeding diet on male KHK ^+/+^ and KHK^-/-^ C57BL/6J mice.

|  | Male KHK <sup>+/+</sup> |  |  | Male KHK <sup>-/-</sup> |  |  |
| --- | --- | --- | --- | --- | --- | --- |
|  | Low Fat | HFHS | HFHD | Low Fat | HFHS | HFHD |
| Insulin (ng/mL) | 1.76±0.36 | 2.92±0.48 | 2.06±0.22 | 1.64±0.25 | 1.20±0.14 | 1.82±0.24 |
| Leptin (pg/mL) | 5974.38±1633.26 | 8062.89±1237.59 | 9557.36±2273.19 | 3640.07±1049.77 | 5737.85±998.88 | 11120.93±2285.07<sup>##</sup> |
| FGF-21 (pg/mL) | 2294.15±260.03 | 1706.17±415.35 | 1501.46±281.91 | 1820.04±524.88 | 2127.62±594.29 | 5560.47±1142.53<sup>##/-</sup> |
| Glucose (mg/dL) | 144.75±8.67 | 184.4±11.36<sup>*</sup> | 177±9.74 | 156.44±6.80 | 210.33±15.03<sup>**</sup> | 205.78±12.07<sup>*</sup> |
| Triglycerides (mg/dL) | 174.02±4.12 | 178.65±10.99 | 178.57±15.77 | 69.93±9.73 | 68.43±4.05 | 76.41±5.96 |
| Cholesterol (mg/dL) | 92.23±12.61 | 151.13±13.77<sup>**</sup> | 151.88±9.54<sup>##</sup> | 85.12±10.77 | 75.95±3.01 | 107.38±11.08 |
| BHB (mmol/L) | 0.18±0.04 | 0.30±0.05 | 0.19±0.05 | 0.14±0.02 | 0.24±0.07 | 0.13±0.02 |
| NEFA (mEQ/L) | 0.54±0.06 | 0.72±0.03 | 0.72±0.04<sup>#</sup> | 0.61±0.05 | 0.36±0.05<sup>*</sup> | 0.53±0.05 |

\**p*<0.05, \*\*/<sup>###</sup> *p*<0.0083. \* Low Fat vs HFHS, <sup>#</sup>Low Fat vs HFHD, <sup>~</sup>HFHS vs HFHD. Data are mean ± SEM. n= 8-10 mice per group. Ordinary one-way ANOVA with Tukey's multiple comparisons test.

We and others previously reported that dietary fructose elongates the intestinal villi, enabling improved nutrient absorption^2,30^. Using H&E-stained duodenal sections, we found that HFHS, but not HFHD, led to a ∼40% elongation of villi in KHK^+/+^ mice but not KHK^-/-^ mice (**Figure 1E, 1F**). Altogether, these findings establish KHK as an essential mediator of fat mass expansion under HFHS but not HFHD feeding.

To directly test whether whole body fructose oxidation requires KHK, we quantified the conversion of labeled substrate into ^13^CO_2_ in age matched male KHK^+^/^+^ and KHK^-^/^-^ mice fed a HFHS diet containing either ^13^C-glucose or ^13^C-fructose. Continuous monitoring in metabolic cages revealed robust oxidation of glucose in the post-prandial state in both genotypes, consistent with substantial glucose utilization (**Figure 1G, Figure S1H**). On the other hand, fructose oxidation showed a striking dependence on KHK, where ^13^CO_2_ production from ^13^C-fructose was robust in KHK^+^/^+^ but nearly absent in KHK^-^/^-^ mice (**Figure 1H, Figure S1G**). We next asked whether pharmacologic inhibition of KHK would phenocopy genetic KHK loss. For this purpose, we used PF-06835919, a small-molecule KHK inhibitor (KHKi) that has been shown in *in vitro* assays to be highly selective, with no observed toxicity in rats or dogs^31^.

Treatment with KHKi substantially reduced fructose-derived ^13^CO_2_ production in KHK^+^/^+^ mice, with minimal effect on glucose-derived ^13^CO_2_ (**Figure 1G, H**). Notably, PF-06835919 had no additional effect in KHK^-^/^-^ mice, suggesting that the drug acts on target. Collectively, these data establish that fructose oxidation during HFHS feeding is highly dependent on KHK activity and can be effectively suppressed by selective KHK inhibition.

### Pharmacological KHK inhibition promotes weight loss by suppressing adiposity and delaying lipid absorption in DIO mice

Pharmacologic inhibition of KHK has previously been shown to prevent fructose-induced hyperlipidemia, hyperinsulinemia, and adipose expansion in rodents^28,32^. Therefore, we examined the effects of PF-06835919 on adiposity during HFHS feeding in obese mice, previously studied in mice and humans with metabolic dysfunction-associated fatty liver disease and type 2 diabetes^28,31,33,34^. Mice were fed a HFHS diet for 15 weeks to induce obesity and then were randomly assigned into three groups for a 4-week intervention: continued HFHS plus vehicle control (HFHS), continued HFHS plus KHKi (HFHS + KHKi), or diet swap to HFHD plus vehicle control (HFHD). Longitudinal body weight monitoring revealed a significant weight loss in the KHKi treated group shortly after treatment initiation (**Figure 2A**). On average, KHKi and HFHD groups lost ∼34% and ∼10% of their body weight by the end of the study, respectively (**Figure 2B**). Analysis of body composition confirmed that the weight loss in the KHKi group was primarily attributed to reductions in fat mass (**Figure 2C**), with lean mass remaining unaffected (**Figure 2D, S3A-C**). These findings were consistent with marked reductions in insulin, triglycerides (TG), leptin, and fibroblast growth factor-21 (FGF-21) levels in KHKi treated mice (**Figure 2E**, **Table 2**), reflecting a broad improvement in metabolic health^32^. Of note, KHKi led to no changes in body weight, lean mass, nor fat mass when given to KHK^-/-^ mice fed HFHS, suggesting an on-target effect (**Figure S3H-J**).

**Figure 2.**
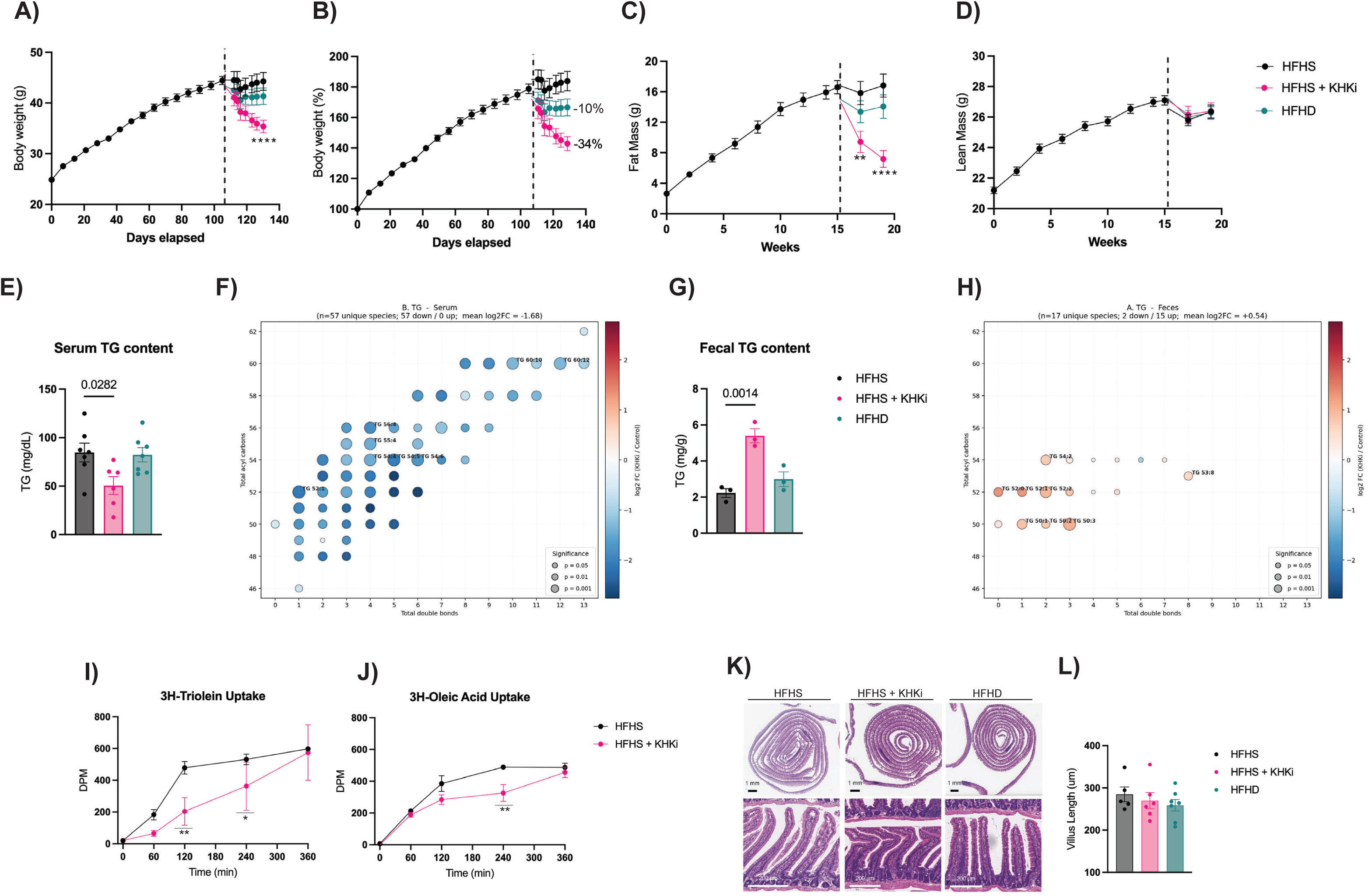
Pharmacological KHK inhibition broadly reduces circulating triglyceride species in serum of DIO mice: Male C57BL/6J mice were fed a high-fat high-sucrose (HFHS, black) diet for 15 weeks to develop a diet-induced obese (DIO) model. After this time, mice were randomized and subdivided in three different groups: continued HFHS (vehicle), HFHS + KHKi (PF-06835919, 50 mg/kg, pink) or high-fat high-dextrose (HFHD + vehicle, teal). **A)** Body weight over time and **B)** Relative weight change over time show significant weight reduction in the KHKi group (n=10 mice per group). **C)** Fat mass and **D)** Lean mass were measured longitudinally by body composition analysis, revealing that fat mass was significantly reduced in KHKi treated mice, while lean mass was intact (n=10 mice per group). Triglyceride (TG) content in **E)** Serum and **G)** Feces were measured at the end of the experiment under the stimulus of different diet exposures (n=3-7 mice per group). **F)** Bubble plot showing the log_2_ fold change (KHKi *vs*. vehicle) of serum TG molecular species in DIO mice. Each bubble represents one TG species, positioned by total acyl carbon number (y-axis) and total number of double bonds (x-axis) (n=3-4 mice per group). **H)** Bubble plot showing the log_2_ fold change (KHKi *vs*. vehicle) of fecal TG molecular species in DIO mice (n=5 mice per group). Selected species are labeled for reference. Serum lipid absorption was measured following oral gavage of radiolabeled lipids. **I)** Serum uptake of [^3^H] triolein over 360 min, and **J)** Serum uptake [^3^H] oleic acid over 360 min (n=2-4 mice per group). **K)** Representative hematoxylin and eosin (H&E)-stained sections of the proximal small intestine show intact villus structure across all groups. Scale bars: black: 1 mm (low magnification); white bar: 200 µm (high magnification). **L)** Quantification of villus length confirms that KHKi treatment does not impair intestinal morphology at the macroscopical level (n=5-7 mice per group). Data is presented as mean ± SEM. *P* values were calculated using either two-way analysis of variance (ANOVA) with Tukey’s multiple comparison test or ordinary one-way ANOVA. \**p*=0.05, \*\**p*=0.001. **** *p*<0.0001. Individual data points are overlaid.

**Table 2:**
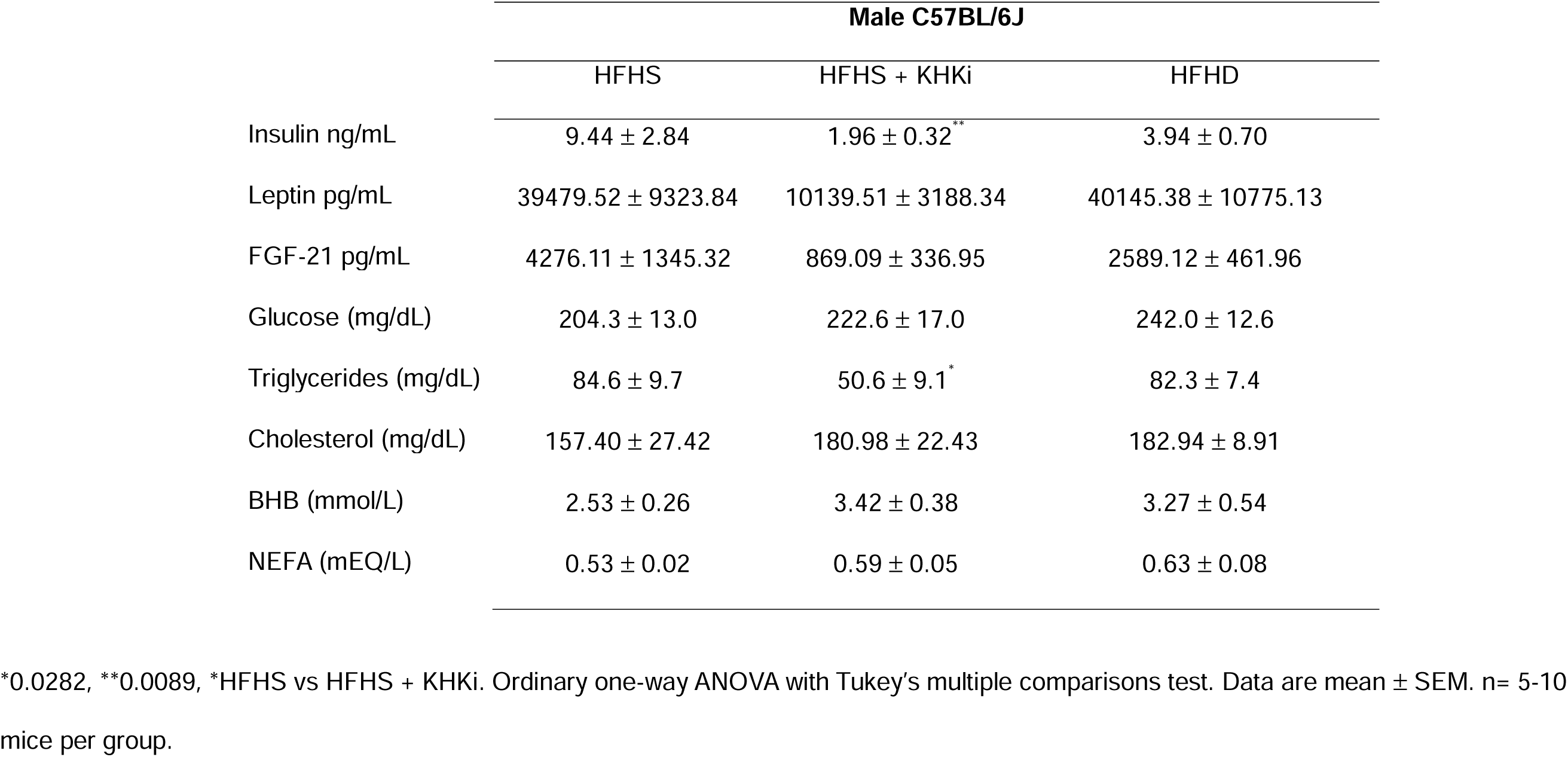
Effects of PF-06835919 intervention under a HFHS diet on male C57BL/6J mice.

|  | Male C57BL/6J |  |  |
| --- | --- | --- | --- |
|  | HFHS | HFHS + KHKi | HFHD |
| Insulin ng/mL | 9.44 ± 2.84 | 1.96 ± 0.32<sup>**</sup> | 3.94 ± 0.70 |
| Leptin pg/mL | 39479.52 ± 9323.84 | 10139.51 ± 3188.34 | 40145.38 ± 10775.13 |
| FGF-21 pg/mL | 4276.11 ± 1345.32 | 869.09 ± 336.95 | 2589.12 ± 461.96 |
| Glucose (mg/dL) | 204.3 ± 13.0 | 222.6 ± 17.0 | 242.0 ± 12.6 |
| Triglycerides (mg/dL) | 84.6 ± 9.7 | 50.6 ± 9.1<sup>*</sup> | 82.3 ± 7.4 |
| Cholesterol (mg/dL) | 157.40 ± 27.42 | 180.98 ± 22.43 | 182.94 ± 8.91 |
| BHB (mmol/L) | 2.53 ± 0.26 | 3.42 ± 0.38 | 3.27 ± 0.54 |
| NEFA (mEQ/L) | 0.53 ± 0.02 | 0.59 ± 0.05 | 0.63 ± 0.08 |

\*0.0282, \*\*0.0089, \*HFHS vs HFHS + KHKi. Ordinary one-way ANOVA with Tukey's multiple comparisons test. Data are mean ± SEM. n= 5-10 mice per group.

In contrast to prior rat studies reporting that pharmacologic KHK inhibition improves metabolic parameters without altering food intake, KHKi treatment in our model significantly reduced food intake, without altering total energy expenditure (**Figure S3D-E**). Cumulative food intake was significantly lower in KHKi-treated mice compared with HFHS controls, most prominently during the dark phase of the light–dark cycle, corresponding to the animals’ peak feeding period. In contrast, mice switched to the HFHD diet consumed the greatest total amount of food, indicating that substitution of sucrose with dextrose substantially alters feeding behavior.

While food intake and energy expenditure are the primary contributors, nutrient absorption is an underrecognized factor that can alter energy balance. KHKi treatment was also associated with reduced caloric absorption relative to both HFHS and HFHD groups (**Figure S3F**), resulting in elevated fecal triglyceride content in KHKi-treated mice, as compared to HFHS or HFHD controls (**Figure 2G**). To confirm whether KHKi affects fat absorption, we performed lipidomics analyses on serum and fecal samples from these mice. Strikingly, DIO mice under the KHKi treatment exhibit a significant reduction in circulating mid-range carbon chain lengths (C50-C54) with low-to-moderate unsaturation (1-5 double bonds) (**Figure 2F, Figure S3L, N**). Furthermore, higher levels of TG content were detected in the fecal samples of the KHKi (**Figure 2E, Figure S3K**). Together, these findings suggest that KHKi reduces adiposity through both decreased food intake and impaired lipid absorption.

To assess whether the serum TG species depleted by KHKi in our mouse model correspond to species elevated by dietary fructose in an independent dataset, we cross-referenced our serum lipidomics against the publicly available in vivo mouse serum lipidomics from Fowle-Grider et al.^34^ In that study, C57BL/6J mice were provided high-fructose corn syrup (HFCS) in drinking water (n=6) or normal water (Control, n=4) for 10 weeks, and serum lipids were profiled by untargeted LC-MS/MS. Of 40 TG species detected in Fowle-Grider et al. and 57 detected in our serum dataset, 33 were present in both. Among the 33 overlapping species, 15 showed convergent directionality: HFCS elevated these species in Fowle-Grider et al. while KHKi depleted them in our dataset. These convergent species were uniformly low-unsaturation TGs (1–7 double bonds; mean 2.9 DB), consistent with de novo lipogenesis-derived oleic acid-containing TGs. Twelve of the 15 convergent species reached statistical significance in our serum dataset (**Figure S3M**). The remaining 18 overlapping species showed divergent directionality: both HFCS and KHKi depleted these species, which were uniformly high-unsaturation TGs (5–13 double bonds; mean 7.6 DB), consistent with dietary PUFA-containing TGs that are displaced by de novo synthesized MUFA-TGs under HFCS feeding and are also reduced by KHKi due to global impairment of intestinal TG export.

Dietary lipids are hydrolyzed and emulsified in the duodenum, which enables fatty acids uptake by the enterocytes^10^. To directly assess whether KHK inhibition impairs the absorption of dietary lipids, we performed metabolic tracing experiments. Specifically, we orally administered triglyceride ([^3^H] triolein) or fatty acid ([^3^H] oleic acid) to fasted obese mice treated with KHKi or vehicle and measured the appearance of radioactivity in the plasma over time, and body tissues at endpoint. The KHKi treatment significantly reduced early intestinal uptake of both tracers, with marked suppression of triolein uptake at 120 minutes and delayed oleic acid uptake at 240 minutes compared with HFHS controls (**Figure 2I, 2J**). Notably, tracer uptake in KHKi-treated mice progressively increased at later time points and partially converged with control levels by 360 minutes, indicating a delay rather than a complete block in lipid absorption. Consistent with delayed lipid absorption, regional analysis of tracer distribution revealed reduced accumulation of labeled oleic acid in the proximal small intestine of KHKi-treated mice compared with other high-fat groups (**Figure S3G**), whereas fatty acid uptake in more distal intestinal segments was comparable across groups. As expected, tracer recovery in the colon was minimal in all conditions; however, KHKi treatment was associated with a trend toward increased lipid recovery in this region, consistent with distal redistribution of unabsorbed dietary fat. In parallel, hepatic accumulation of labeled lipids tended to be lower in KHKi-treated mice relative to both HFHS and HFHD controls, suggesting reduced systemic delivery of intestine-derived lipids at this timepoint. These data demonstrate that KHK inhibition slows the rate of postprandial dietary lipid appearance, consistent with altered intestinal lipid handling and delayed chylomicron-mediated lipid delivery.

Because KHK has been shown to enable intestinal villus elongation following HFHS exposure (**Figure 1E, 1F**), we hypothesized that the delayed lipid absorption observed with KHK inhibition might reflect a reduction in absorptive surface area due to villus shortening, potentially in a region-specific manner within the proximal intestine. To address this question, we quantified villus length by histologic analysis of H&E–stained sections. Contrary to our hypothesis, villus length was not reduced in KHKi-treated or HFHD-fed mice compared with HFHS controls in either the duodenum (**Figure 2K–L**) or more distal intestinal segments. These findings indicate that delayed lipid absorption with KHK inhibition occurs independently of gross changes in villus length.

### Fructolysis and chylomicron assembly programs are spatially coordinated in the proximal intestine

We next considered whether fructose metabolism influences dietary lipid transport at the level of enterocyte lipid handling. In both mouse and human small intestine, enterocytes undergo extensive transcriptional remodeling along two orthogonal spatial axes: the proximal-to-distal axis, which reflects regional specialization for nutrient absorption, and the crypt-to-villus axis (denoted as V1 to V6), which reflects enterocyte maturation state^8,9,35^. We re-examined publicly available single-cell RNA-sequencing data from enterocytes along these axes to identify the expression patterns of genes encoding fructolysis enzymes (Khk, Slc2a5, Aldob), chylomicron assembly factors (Mttp, Apob, Mogat2, Dgat1, Dgat2, Cideb), lipid droplet and fatty acid uptake markers (Plin2, Cd36, Fabp2), and apolipoproteins (Apoa4, Apoa1).

Along the proximal-to-distal axis, fructolysis and chylomicron assembly genes are co-enriched in the proximal small intestine in both mouse and human (**Figure S4A/B**). In mouse enterocytes, Khk and Slc2a5 were enriched in proximal intestinal segments, paralleling expression of chylomicron assembly genes including Mttp and Mogat2. Across the proximal-to-distal axis, Khk and Mttp expression were strongly correlated (Pearson r = 0.939, p < 0.0001), consistent with coordinated spatial organization of fructolysis and lipid export programs along this axis. An equivalent pattern was observed in human enterocytes, where KHK, SLC2A5, MTTP, MOGAT2, and DGAT1 all peaked in the most proximal segment pairs, with CD36 again showing the steepest gradient (15.5 proximal enrichment) and KHK–MTTP co-expression preserved (Pearson r = 0.932, p < 0.0001). These data localize fructose catabolism and chylomicron assembly programs to the proximal small intestine.

We explored the gene expression changes in the crypt-to-tip axis at a single proximal-to-distal position in the jejunum using data from Moor *et al.* (2018)(**Figure S4C/D**).^35^ Fructolysis genes display a mid-villus-biased expression pattern, peaking at the mid villus (V2–V3) and declining toward both the crypt and the villus tip. For example, Khk mean expression rises from 0.88 in the Crypt to a peak of 2.08 at V2, then declines to 0.35 at V6 (tip). Slc2a5 follows a similar pattern, peaking at V3, and Aldob is more broadly expressed but also peaks at V2. Chylomicron assembly genes are broadly tip-biased, with expression rising from the crypt and plateauing across V2–V5. Apolipoproteins show monotonically increasing expression from crypt to tip, consistent with their role in chylomicron maturation at the villus surface. A similar crypt-to-tip organization of fructolysis and lipid export genes was observed in human intestinal tissue by spatial transcriptomics^8^. The spatial offset between fructolysis (mid) and chylomicron assembly (tip) along the crypt-to-tip axis is consistent with a model in which fructose catabolism via KHK in mid-villus enterocytes provides substrate or regulatory signals that promote downstream lipid packaging and chylomicron export in more apical cells.

### KHK inhibition reprograms enterocyte lipid trafficking and chylomicron assembly

Together, these spatial transcriptomic analyses suggested that fructose metabolism may directly regulate intracellular enterocyte lipid trafficking. Because cytoplasmic lipid droplets (CLDs) function as transient intermediates in chylomicron assembly and dietary lipid export^36^, we next tested whether pharmacologic inhibition of KHK alters CLD formation and regional lipid handling in vivo. CLDs are particularly abundant in the proximal small intestine and serve as a buffering compartment during periods of high lipid flux, thereby shaping postprandial lipid delivery to the circulation^37^. To determine whether KHK inhibition alters this intracellular lipid trafficking step, we examined the intestine by transmission electron microscopy (TEM) following an oral gavage of olive oil. Mice were exposed to 10 days of water or HFCS to allow for any potential enterocyte reprogramming induced by fructose exposure, and half the mice from each group were co-administered KHKi daily to block the effects of fructose. The resulting tissues were fixed and stained with 2% osmium tetroxide, which reacts with unsaturated fatty acids in the CLDs to increase the electron density on TEM^38^.

In the vehicle control group (H_2_O+ DMSO), CLDs were readily observed in mid-villus enterocytes as electron-dense cytoplasmic structures (red arrows), with larger droplets (i.e. >15 µm^2^) present in the jejunum (47.68±5.65) (p<0.0001, Welch’s test) compared with the duodenum (20.99±1.91) (**Figure 3A, 3D**). These findings are consistent with spatial transcriptomic analyses identifying mid-villus enterocytes as a major site of fructolytic gene expression in both mouse and human intestine (**Figure S4 C, D**). HFCS exposure resulted in marked enlargement of CLDs in the duodenum accompanied by smaller and less electron-dense CLDs in the jejunum and ileum, consistent with a proximal redistribution of lipid handling (**Figure 3B, 3E, 3H**). These regional patterns were profoundly altered by KHK inhibition. In mice not exposed to HFCS, KHKi treatment nearly abolished CLD formation across all intestinal segments examined (**Figure 3A, 3D, 3G**). In HFCS-treated mice, KHKi markedly reduced CLD size and density in the duodenum while increasing the presence of larger, less dense CLDs in the jejunum and ileum, consistent with a shift in lipid processing toward more distal intestinal segments (**Figure 3C, 3F, 3I**). Together, these findings suggest that fructose metabolism promotes proximal enterocyte lipid storage and chylomicron processing, whereas KHK inhibition delays and redistributes intestinal lipid trafficking toward more distal segments.

**Figure 3.**
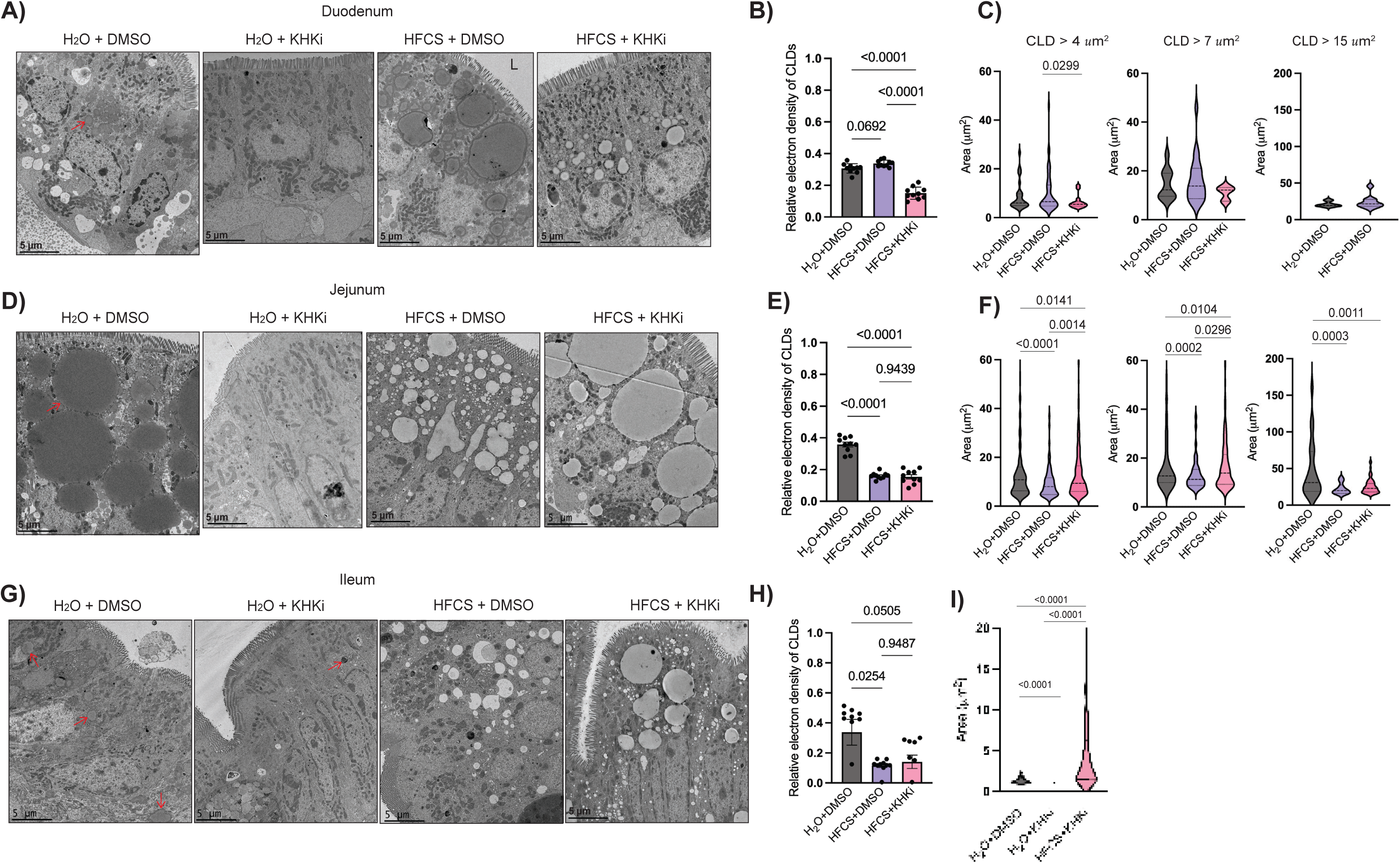
PF-06835919 reduces cytoplasmic lipid droplet formation in different sections of the small intestine. Representative electron microscopy (EM) images of absorptive enterocytes and cytoplasmic lipid droplets (CLDs) from each region of the small intestine in response to dietary fat. Mice were fasted overnight, treated with either water or High Fructose Corn Syrup (HFCS) and with or without PF-06835919, after an hour, a 200 μl olive oil oral gavage was administrated, and 2 h later samples were collected (n=3 mice per group). CLDs are detected as round organelles (red arrowhead) inside of the enterocytes from the **A)** Duodenum, **D)** Jejunum and **G)** Ileum regions of the small intestine. ‘L’ indicates lumen. Scale bar is 5 μm. Electron density analysis in the CLDs of ten intact cells per mouse in the **B)** Duodenum and **E)** Jejunum and **H)** Ileum regions reveal changes in lipid composition under the different treatments. Total CLD area per cell of all cells analyzed per mouse and size are represented as violin plots, revealing differences across experimental conditions in the **C)** Duodenum, **F)** Jejunum and **I)** Ileum region. Water + DMSO (H_2_O + DMSO, black), water + KHKi (H_2_O + KHKi, red), HFCS + DMSO (purple) and HFCS + KHKi (pink), n=3 mice per group. H_2_O + KHKi group is not included as KHKi almost completely prevented the formation of CLDs that the script could detect.

Because KHK inhibition redistributed CLD morphology across intestinal regions, we next asked whether fructose metabolism also regulates the spatial organization of chylomicron assembly machinery. Lipids stored within enterocyte CLDs are mobilized through regulated lipolysis and CLD–endoplasmic reticulum coupling, supplying re-esterified triglycerides to the microsomal triglyceride transfer protein (MTTP)–apolipoprotein B (apoB) complex for chylomicron lipidation and secretion^36,39^. Consistent with this role, genetic ablation of MTTP in mice blocks intestinal triglyceride absorption and chylomicron secretion, and is associated with reduced ApoB export from enterocytes^40^. Given that MTTP can function as a rate-limiting component of chylomicron assembly and thus influence delivery of dietary lipids to peripheral tissues, we next examined whether blocking fructose metabolism alters the regional expression of chylomicron assembly machinery. To this end, we performed RNAscope in situ hybridization for *Mttp* across the small intestine under the indicated dietary and pharmacologic conditions.

In the proximal duodenum, overall *Mttp* transcript abundance was not significantly altered across dietary interventions (**Figure 4A–C**). However, marked differences in spatial distribution were observed along the crypt-villus axis, with *Mttp* expression enriched near the crypt region relative to the villus tip under LF and HFHD conditions. In contrast, HFHS feeding was associated with a more uniform distribution of *Mttp* expression along the proximal and distal duodenum (**Figure 4D-F**), whereas KHKi treatment resulted in a relative shift of *Mttp* expression toward more distal intestinal segments (**Figure 3G–I**). Collectively, these findings suggest that fructose metabolism spatially organizes intestinal chylomicron assembly machinery to favor proximal lipid processing, whereas KHK inhibition shifts this program toward more distal intestinal regions.

**Figure 4.**
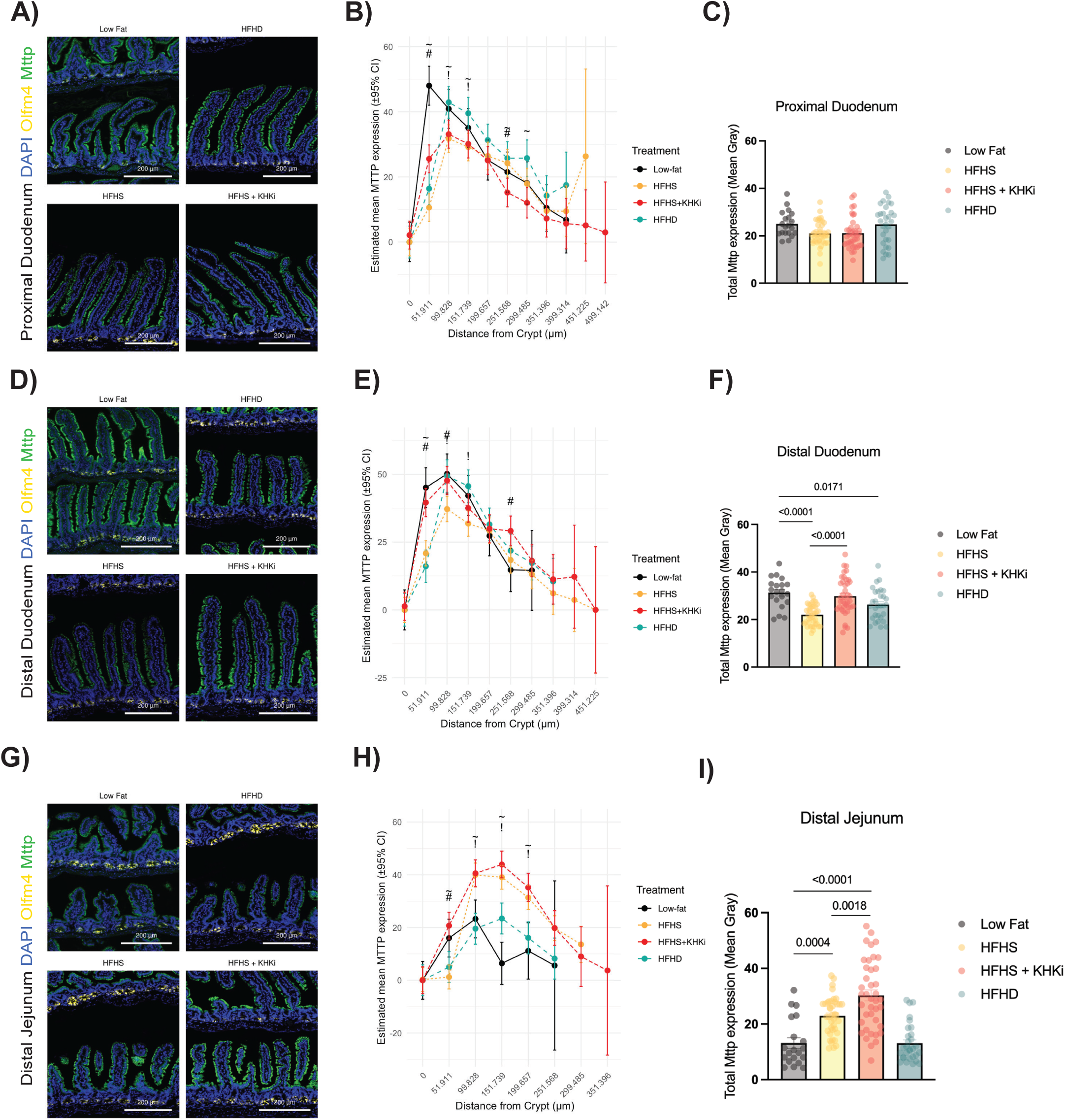
Ketohexokinase inhibition stimulates distal compensatory effect in the small intestine under the presence of fructose: RNAscope *in situ* hybridization of *Mttp* in the **A)** Proximal duodenum, **D)** Distal duodenum and **G)** Distal jejunum sections under the exposure of different diet interventions. *Mttp* expression in green (Opal 520), *Olfm4* (Opal 570) in yellow and DAPI in blue. Scale bar: 200 µm. Quantification of *Mttp* expression across the villus length (from crypt to tip) of 10 villi per mouse (n=3-4 mice per group) in the **B)** Proximal duodenum, **E)** Distal duodenum and **H)** Distal jejunum section. Total *Mttp* expression in the **C)** Proximal duodenum, **F)** Distal duodenum, and **I)** Distal jejunum area under different diets. Low fat (grey), HFHS (High-fat high-sucrose; yellow), HFHS+ KHKi (High-fat high-sucrose + PF-06835919; red), HFHD (High-fat high-dextrose, teal). *P* values were calculated using two-way analysis of variance (ANOVA) with Tukey’s multiple comparison test or one-way ANOVA with Tukey’s multiple comparison or Welch’s test. Data are presented as mean ± SEM; individual data points are overlaid. ^#^ HFHS: KHKi, ^^^ KHKi: Low fat, ^∼^HFHD: KHKi ^!^ HFHD: HFHS ^+^HFHS: Low Fat ^-^ HFHD:Low Fat

### KHK inhibition enhances incretin-based weight loss independently of appetite suppression

The finding that KHK inhibition reduces intestinal lipid absorption through an enterocyte-intrinsic mechanism raised the possibility that it might complement incretin-based therapies, which promote weight loss primarily through central suppression of appetite^41^. We therefore asked whether pharmacologic blockade of fructose metabolism could augment the efficacy of semaglutide, a glucagon-like peptide 1 (GLP-1) receptor agonist^42–44^, by targeting a distinct and non-redundant step in energy harvest.

To test this question, we randomized obese mice to receive vehicle treatment, semaglutide alone, or the combination of semaglutide and KHKi for 4 weeks while continuing HFHS feeding. Consistent with prior reports^41,45^, semaglutide decreased body weight by 30% compared to vehicle treated mice (**Figure 5A, 5B**). Notably, the addition of KHKi to semaglutide produced a greater reduction of approximately 40%. Body composition analysis showed that semaglutide, alone and in combination, exhibited similar reductions in lean mass when compared to vehicle control group; whereas the reduction in fat mass tended to be lower in KHKi treated mice (**Figure 5C, 5D**). The reduction in total body fat mass was reflected in the masses of the inguinal and gonadal white adipose depots, and the reduction in lean mass was reflected by the masses of the liver and gastrocnemius (**Figure S4A-F**). Semaglutide alone or in combination with KHKi also significantly reduced serum levels of insulin, leptin, glucose, TG, and cholesterol (**Table 3**), reflecting improvement in metabolic health.

**Figure 5.**
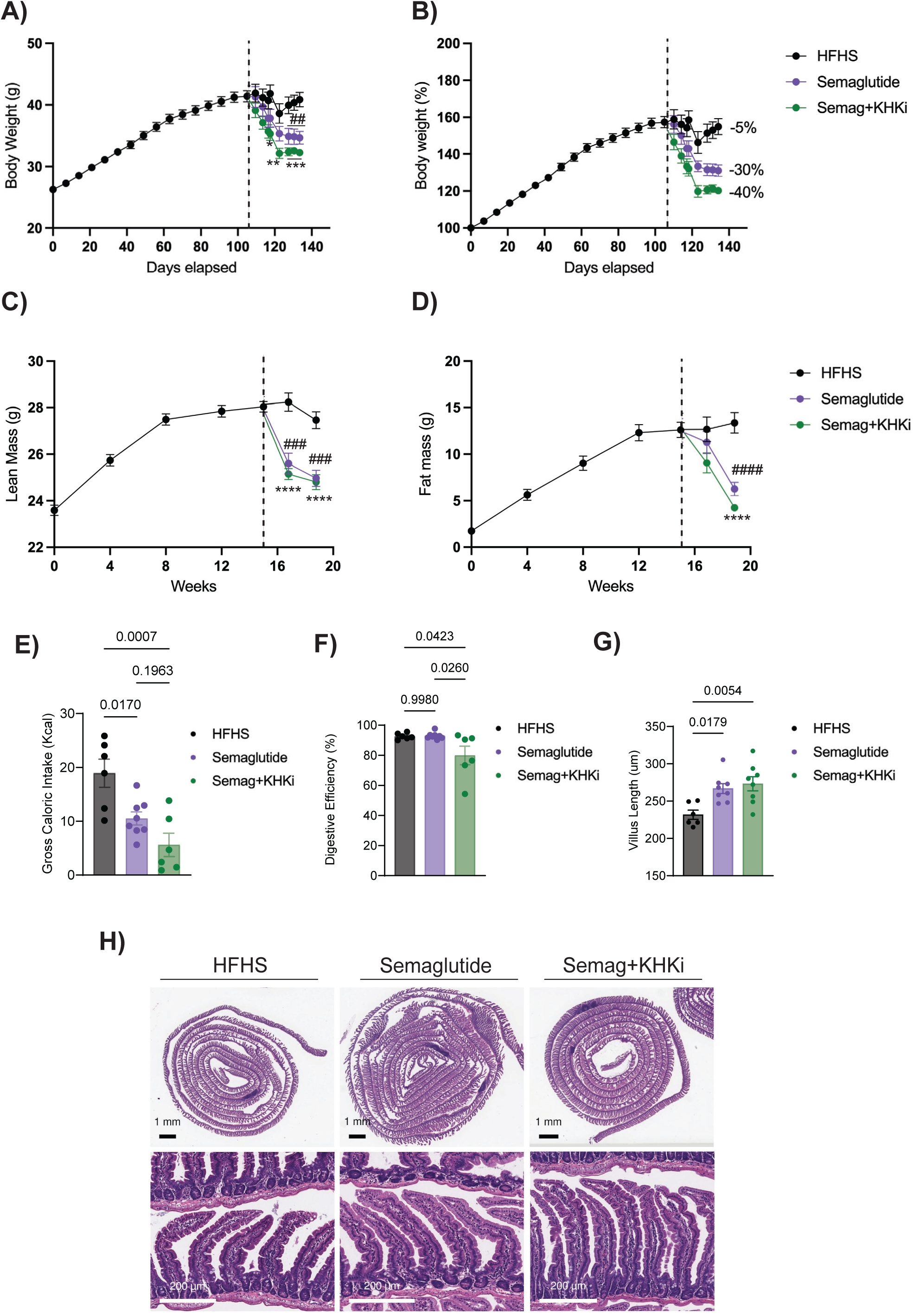
Combination of semaglutide and PF-06835919 improves fat loss in mice fed under a high-fat high-sucrose diet. Male C57BL/6J mice were fed a high-fat high-sucrose (HFHS; black) diet for 15 weeks. After this time, mice were randomized and subdivided in three different groups: continued HFHS (vehicle control, black), continue HFHS + semaglutide (30 nmol/kg; purple) or continue HFHS + semaglutide + KHKi (PF-06835919; 50 mg/kg; green). **A)** Body weight over time and **B)** Relative weight change (%) over time (n= 9-10 mice per group). **C)** Lean mass and **D)** Fat mass over time. **E)** Gross caloric intake (kcal) and **F)** Digestive Efficiency (%). **G)** Proximal villus length after 4 weeks of semaglutide or combination of semaglutide and PF-06835919 exposure. **H)** Representative haematoxylin and eosin (H&E)-stained proximal intestine from DIO mice under the exposure of different treatments. Black bar: 1 mm; white bar: 200 µm. Data are presented as mean ± SEM with individual data points overlaid. *P* values were calculated using two-way analysis of variance (ANOVA) with Tukey’s multiple comparison test. *^/#^ *p*<0.05, ^**/##^ *p*<0.001, ^***/###^ *p*<0.0001.

**Table 3:** Effects of semaglutide and PF-06835919 interventions under a HFHS diet on DIO male C57BL/6J mice.

| DIO Male C57BL/6J |  |  |  |
| --- | --- | --- | --- |
|  | HFHS | Semaglutide | Semaglutide + KHKi |
| Insulin ng/mL | 5.18 ± 0.81 | 1.92±0.23<sup>***</sup> | 1.01±0.22<sup>****</sup> |
| Leptin pg/mL | 28359.16 ± 4428.92 | 6044.20±1143.69<sup>****</sup> | 1674.46±207.36<sup>****/~</sup> |
| FGF-21 pg/mL | 2162.53 ± 409.37 | 1657.35 ± 398.78 | 2101.17 ± 241.04 |
| Glucose (mg/dL) | 203.33 ± 6.71 | 176.50 ± 9.59<sup>*</sup> | 123.56 ± 5.09<sup>~###</sup> |
| Triglycerides (mg/dL) | 213.63 ± 8.77 | 175.26 ± 3.90<sup>*</sup> | 154.19 ± 14.26<sup>~</sup> |
| Cholesterol (mg/dL) | 150.06 ± 7.61 | 102.47 ± 7.38<sup>****</sup> | 87.48 ± 2.97<sup>~</sup> |
| BHB (mmol/L) | 0.11 ± 0.02 | 0.12 ± 0.02 | 0.16 ± 0.06 |
| NEFA (mEQ/L) | 0.18 ± 0.03 | 0.23 ± 0.05 | 0.27 ± 0.07 |

<sup>\*\*\*/###</sup> $p=0.001$ , <sup>\*\*\*\*/####/~</sup> $p<0.001$ . \*HFHS vs Semaglutide, ~ HFHS vs Semaglutide + KHKi, # Semaglutide vs Semaglutide + KHKi. Ordinary one-way ANOVA with Tukey's multiple comparisons test. Data are mean ± SEM. n= 9-10 mice per group.

To determine whether the greater weight loss observed with combined semaglutide and KHKi treatment could be explained by differences in energy balance, we assessed the gross caloric intake and digestive efficiency in singly housed mice over a 48-hour period using metabolic chambers. Semaglutide treatment significantly reduced gross caloric intake compared with HFHS controls, consistent with appetite suppression. There was trend for caloric intake to be further reduced with the KHKi relative to semaglutide alone, but this difference was not statistically significant (**Figure 5E**). Digestive efficiency provides an estimate of how effectively ingested energy is absorbed by the host and is calculated as the ratio of absorbed energy to total energy intake. Using this metric, digestive efficiency was similar between HFHS- and semaglutide-treated mice, whereas combined semaglutide and KHKi treatment resulted in a significant reduction in digestive efficiency, indicating impaired nutrient absorption (**Figure 5F**). Total energy expenditure (TEE) was similar between treated animals versus control group (**Figure S4G**). These findings indicate that the additional ∼10% weight loss achieved with combined semaglutide and KHKi treatment is associated with both reduced food intake and a further decrease in nutrient absorption efficiency, rather than changes in energy expenditure.

We next evaluated the effects of semaglutide and the combination therapy on intestinal villus structure. Surprisingly, the semaglutide treated mice displayed longer and more dense villi in the proximal intestine, which was not altered with KHKi treatment (**Figure 5G, 5H**). These data suggest that the reduction in absorption efficiency with KHK inhibition occurs independently of gross changes in villus length.

Despite robust initial weight loss, discontinuation of GLP-1 receptor agonists such as semaglutide is frequently followed by rapid weight regain^24,46^, highlighting a critical unmet need for strategies that support long-term weight maintenance. To determine whether inhibition of fructose metabolism modifies this response, we performed a semaglutide withdrawal experiment in which obese mice previously treated with semaglutide alone or in combination with KHKi were monitored after cessation of GLP-1 therapy. Once again, semaglutide plus KHKi resulted in more body weight and fat mass loss than semaglutide alone during the initial weight loss phase (**Figure 6A, 6B, Figure S5**). Following withdrawal, mice previously treated with semaglutide alone rapidly regained body weight and adiposity, reaching levels comparable to HFHS controls. In contrast, mice that continued KHKi after semaglutide withdrawal regained weight more slowly and completed the experiment with approximately 17% lower body weight and 20% lower fat mass (**Figure 6B**). These results suggest that the inhibition of KHK can improve weight maintenance following withdrawal of semaglutide.

**Figure 6.**
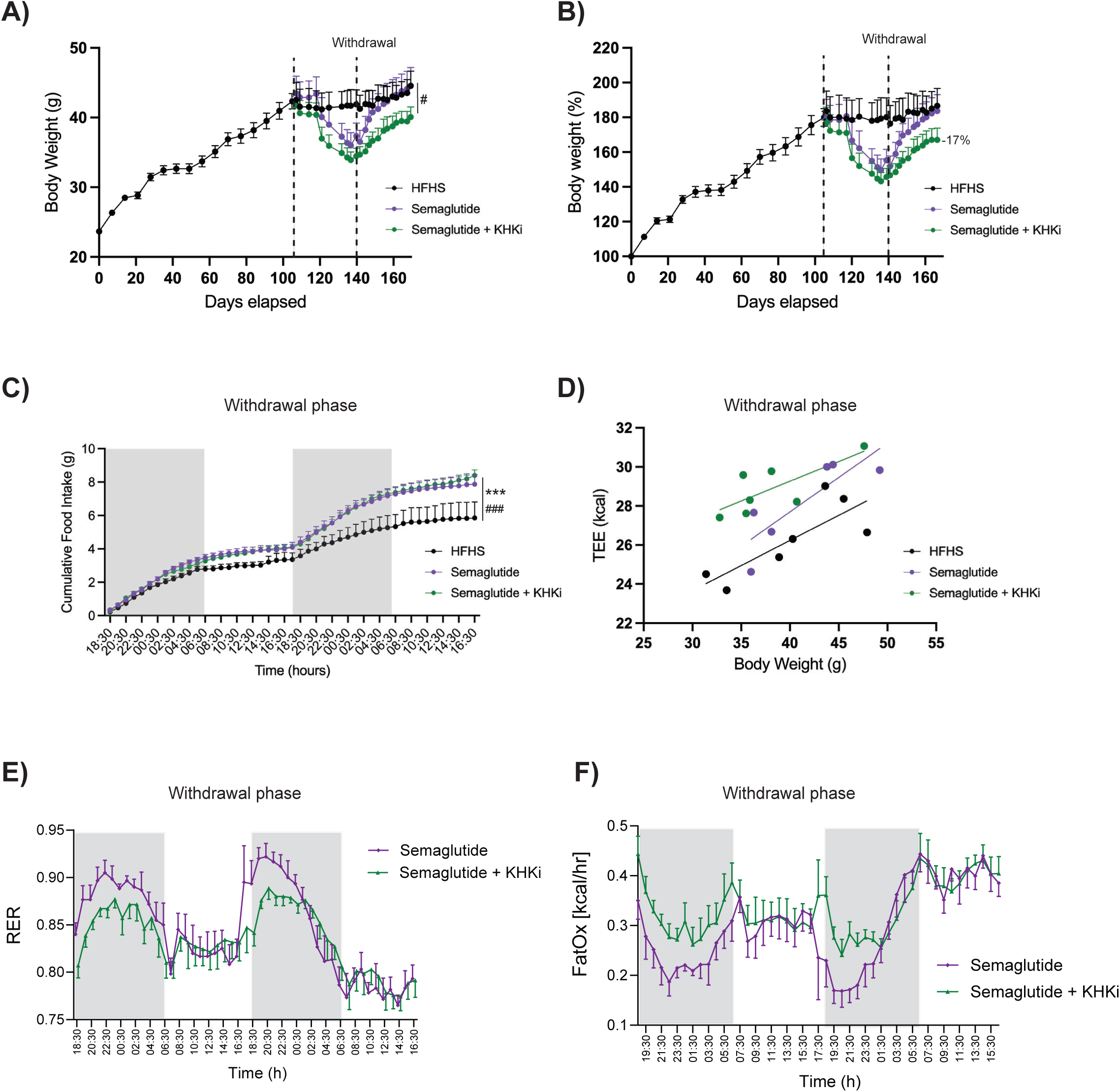
KHKi prevents weight regain independently of food intake and energy expenditure. Male C57BL/6J mice were fed a high-fat high-sucrose (HFHS; black) diet for 15 weeks. After this time, mice were randomized and subdivided in three different groups: continued HFHS + vehicle control, continue HFHS + semaglutide (30 nmol/kg; purple) or continue HFHS + semaglutide + KHKi (PF-06835919; 50 mg/kg; green). At week 20, semaglutide was withdrawn from all experimental groups. **A)** Body weight over time and **B)** Relative weight change (%) over time (n= 6-7 mice per group). **C)** Cumulative food intake during drug withdrawal phase. **D)** Linear regression analyses of total energy expenditure (TEE, kcal) versus body weight (g) over the final 48 hours in metabolic cages during the withdrawal phase. **E)** During the withdrawal experiment, the respiratory exchange ratio (RER) is lower in the continued KHKi group over a period of 48 hours. **F)** High levels of fat oxidation (kcal/hour) are increased in the continued KHKi group versus the semaglutide group. Data are presented as mean ± SEM. *P* values were calculated using two-way analysis of variance (ANOVA) with Tukey’s multiple comparison test. ^#^*p*=0.05, ^***/###^*p*<0.0001. ^#^(HFHS vs Semaglutide+KHKi), *(HFHS vs Semaglutide).

To assess whether differences in energy balance contributed to this effect, we examined food intake and TEE during the withdrawal phase. Food intake increased abruptly after semaglutide discontinuation in both treatment groups, with no significant differences between mice with or without KHKi (**Figure 6C**). When examining TEE in relationship to body weight, we found that the mice maintained on KHKi exhibited higher TEE relative to the other groups (**Figure 6D**). Furthermore, the mice that continued KHKi had a lower RER compared with those previously on semaglutide, particularly during the dark phase, indicating a relative shift toward increased lipid oxidation (**Figure 6E**). Indeed, fat oxidation rates were higher in the KHKi treated group compared with semaglutide alone, despite similar food intake during the withdrawal phase (**Figure 6F**). These data suggest that continued KHK inhibition following GLP-1 withdrawal alters substrate partitioning toward greater reliance on lipid oxidation.

## Discussion

Fructose is a core constituent of high-fat dietary patterns, including Western-style and ultra-processed diets that are strongly associated with obesity and metabolic disease^47,48^. Prior genetic, pharmacologic, and dietary studies in rodents position fructose metabolism as a key contributor to weight gain and metabolic dysfunction arising from these diets, acting through mechanisms that extend beyond caloric load alone^7,12,13,17,18,27,28,49–51^. In this study, we controlled dietary sugar content by replacing sucrose with dextrose, which resulted in less weight gain in lean mice and more weight loss when the obese mice were switched from sucrose to dextrose. These results underscore the specific contribution of fructose metabolism rather than carbohydrate calories per se. Consistent with this interpretation, we also found that pharmacologic inhibition of fructose metabolism promoted weight loss in obese mice both as monotherapy and in combination with semaglutide, indicating that targeting fructose metabolism reduces adiposity independently of and additive to appetite-suppressive therapies.

Our work confirms previous reports that fructose promotes weight gain through a KHK-dependent mechanism^2,7,14,17,18,51^. Mice with whole-body or intestinal-specific KHK knockout are protected from obesity when fed diets rich in fructose-containing sugars. Our data indicate that fructose metabolism via KHK promotes fat accumulation through a dual mechanism, involving a transient increase in food intake during diet initiation and a sustained reprogramming of intestinal lipid metabolism that enhances energy harvest. These findings are consistent with studies showing that intestinal KHK influences feeding behavior and can modify the surface area of intestinal lacteals during high-fructose exposure^7,51^. Together, these data reinforce the importance of the small intestine as an active metabolic gatekeeper for fructose^12–14,28,51,52^.

We and others have reported that dietary fructose elongates the intestinal villi^2,30^. Therefore, we hypothesized that the anti-obesogenic effects of KHK inhibition would be associated with reversal of villus elongation, and a subsequent reduction in nutrient absorption. To our surprise, the treatment had no effect on villus length. However, lipid absorption and energy harvest were significantly reduced with KHKi treatment. These results led us to conclude that the metabolic benefits of KHK inhibition were more likely to reflect changes in intestinal lipid handling at the epithelial cell level, rather than changes in intestinal morphology.

The impact of fructose on intestinal lipid metabolism was visualized by tracking CLDs using TEM. CLDs act as transient, dynamic storage for dietary TGs before they are packaged into chylomicrons and absorbed^36^. Fructose led to regional changes in CLD size and density. In the duodenum, fructose promoted large, electron dense CLDs in the duodenum, which was prevented by KHKi. In the jejunum, fructose led to smaller and less dense CLDs, suggesting a shift in lipid absorption from the jejunum to the duodenum. These results are conceptually similar to those observed with MGAT2 deficiency, which changes the spatiotemporal distribution of fat absorption by reducing lipid absorption in the proximal small intestine and delaying it to more distal segments^53^. In our study, the electron density of CLDs is a reflection of unsaturated fat content^38^. We found that KHKi enriches the feces with unsaturated fats while completely depleting them in the serum, especially TG species. These findings suggest that fructose metabolism may regulate the intestinal handling and systemic availability of unsaturated fatty acids. In this context, ELOVL6 in epithelial cells could play an important role. ELOVL6 catalyzes the elongation of long chain fatty acids and monounsaturated fatty acids with chain lengths from 12 to 16 carbons. *Elovl6^−/−^* mice accumulate palmitic (C16:0) and palmitoleic (C16:1, *n-*7) fatty acids and contain significantly less stearic (C18:0) and oleic (C18:1, *n-*9) acids^54^. These data raise the possibility that fructose metabolism influences oleate availability in enterocytes. Interestingly, Fowle-Grider *et al*. found that fructose feeding increased fasting levels of LPC 18:1 more than sevenfold^34^, and KHKi treatment decreased these levels in the serum^34^.

The small intestine is organized into discrete, functional domains spanning the proximal-distal and crypt-villus axes, each optimized for specific aspects of nutrient processing^8,9,35,55^. These domains exhibit diet-responsive plasticity rather than fixed absorptive roles. In this study, we observed one such example. *Mttp* expression, a marker of chylomicron synthesis, became more uniformly expressed in duodenal epithelial cells along the crypt-villus axis with HFHS feeding, and KHKi treatment resulted in a relative shift of *Mttp* expression toward more distal intestinal segments. We suspect that the changes in *Mttp* expression is controlled by ChREBP. In the liver, fructose activates ChREBP in a KHK-dependent manner, and ChREBP positively modulates *Mttp* expression and lipid export capacity^14,56,57^. Together, these data suggest that KHKi prevents the absorption of unsaturated fatty acids by blocking epithelial lipid metabolism, shifting lipid absorption to more distal regions of the intestine, reducing circulating TGs, and increasing fecal TG excretion.

The redistribution of dietary lipid processing toward distal intestinal segments observed with KHK inhibition has a physiologically important consequence: unabsorbed fat reaching the ileum potently stimulates enteroendocrine L-cells, the primary source of GLP-1 in the gut^58^. L-cell density is highest in the distal ileum and colon^59^, and GLP-1 secretion is driven by luminal lipids, particularly long-chain unsaturated fatty acids^60,61^. Under normal physiology, dietary fat is nearly completely absorbed in the proximal intestine, limiting distal L-cell stimulation. However, the proximal-to-distal shift in lipid trafficking induced by KHK inhibition would be expected to increase luminal fat delivery to the ileum, thereby augmenting endogenous GLP-1 release, which can impact central appetite signaling.

Incretin-based therapies have revolutionized the treatment of obesity; however, long-term compliance rates are low and discontinuation leads to rapid weight regain^62^. Pharmacological strategies targeting two or more receptor systems are gaining attraction for weight loss and weight maintenance. For example, Morningstar et al^63^ used a cannabinoid receptor type 1 inverse agonist to improve weight loss with semaglutide. Here, we demonstrate that inhibition of fructose metabolism improves GLP-1–mediated weight loss, independently of sustained changes in food intake or intestinal villus architecture. The combination of semaglutide and KHKi reduced digestive efficiency, suggestive of changes to intestinal epithelial metabolism as observed when KHKi was given as monotherapy.

Overall, our data support a model in which intestinal fructose metabolism via KHK contributes to enhanced dietary lipid absorption and fat accumulation under HFHS conditions. Pharmacological inhibition of fructose metabolism redistributes lipid handling towards distal segments, which reduces digestive efficiency, contributes to preferential fat mass loss. Therefore, KHK inhibition represents a promising therapeutic strategy for obesity and metabolic disease management, particularly in the context of western diets rich in fat and sugar.

## Limitations of the study

- The present study focused on the intestinal epithelium as the primary site of KHK-dependent lipid handling. However, hepatic fructose metabolism is also substantially altered by KHK inhibition, and prior work has established that KHK-dependent lipid accumulation in the liver contributes to metabolic dysfunction, including steatosis and dyslipidemia^48^.
- KHKi was studied in the presence of HFHS diets. It may not be effective if all fructose is removed from the diet, which was encouraged in clinical studies of PF-06835919 for Metabolic Dysfunction-Associated Fatty Liver Disease (MAFLD).
- PF-06835919 may be absorbed in the proximal intestine and therefore not directly reach distal intestinal epithelial segments directly.
- Fructose at high doses has been shown to alter the gut microbiome and promote gut-barrier deterioration^51,64,65^. Since this effect has been previously established, our focus in this study was the effects on the intestinal epithelium.

## Author Contributions

Conceptualization, C.E.E., M.A., M.D.G., Methodology, C.E.E., M.A., J.G., S.L.S., I.N., L.K.D., C.L., T.A., O.S., T.P., C.P., J.K., P.M., A.E., R.K.Z., D.E.C., Formal Analysis, C.E.E., M.A., J.G., S.L.S., L.K.D., C.L., C.P., M.D.G., Investigation, C.E.E., M.A., J.G., S.L.S., I.N., L.K.D., C.L., O.S., C.P., A.E., P.M., J.K., R.K.Z., L.C.C., D.E.C., M.D.G., Resources, L.K.D., R.K.Z., M.D.G., Data Curation, C.E.E., M.A., J.G., S.L.S., L.K.D., C.L., C.P., M.D.G., Writing– Original Draft, C.E.E., M.A., M.D.G., Writing–Review & Editing: C.E.E., M.A., J.G., S.L.S., I.N., L.K.D., C.L., T.A., O.S., T.P., C.P., A.E., P.M., J.K., R.K.Z., L.C.C., D.E.C., M.D.G. Supervision, L.C.C., M.D.G., Funding Acquisition, M.D.G.

## Acknowledgments

We thank NYU Langone’s Experimental Pathology Research Laboratory (RRID: SCR_017934) and NYULH DART Microscopy Laboratory (partially funded by NYU Cancer Center Support Grant NIH.NCI P30CA016087), Alice Liang, Joseph Sall and Jason Liang for their consultation and assistance with performing the EM work. The Zeiss Gemini300 SEM was purchased with the support of NIH S10OD019974. We acknowledge NYU Langone Health’s Metabolomics Laboratory for its help in acquiring and analyzing the lipidomics data. We would like to thank Dr. Mahmood Hussain for his insightful feedback during the design of the radioactivity experiments.

## Lead contact

Requests for further information and resources should be directed to and will be fulfilled by the lead contact, Marcus D. Goncalves.

---

**Supplementary Figure 1.**
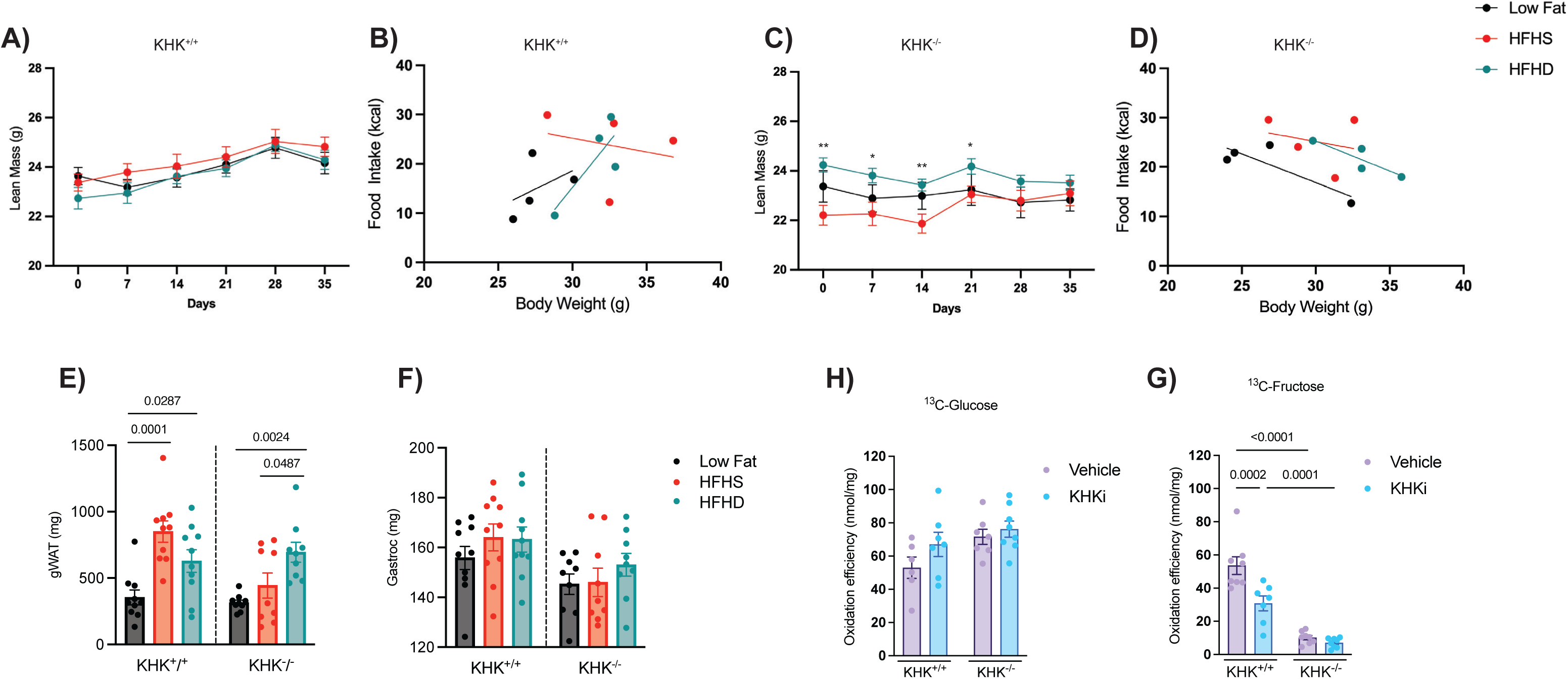
Dietary sucrose promotes adiposity without changing lean mass: **A)** Lean mass over time in KHK^+/+^ animals exposed to different diet interventions over a period of 5 weeks. Low fat (black), high-fat high-sucrose (HFHS, red) or high-fat high-dextrose (HFHD, teal) diets (n= 10 mice per group). **B)** Linear regression analyses of average food intake (kcal) versus body weight over the final 48 hours in metabolic cages from the KHK^+/+^ mice (n= 4 mice per group). **C)** Lean mass over time in KHK^-/-^ mice exposed to different diet interventions over a period of 5 weeks (n= 9 mice per group), *(HFHS vs HFHD), \**p*=0.0264, ** *p*= 0.0045. **D)** Linear regression analyses of average food intake (kcal) versus body weight over the final 48 hours in metabolic cages from the KHK^-/-^ mice (n= 4 mice per group). **E)** After five weeks of different diet exposure, euthanasia was performed, and tissues such as gonadal white adipose tissue (gWAT), and **F)** Gastrocnemius (gastroc), were collected and weighed (n= 9-10 mice per group). Bar graphs showing cumulative **H)**^13^C-glucose and **G)** ^13^C-fructose oxidation efficiency (nmol/mg) in KHK^+^/^+^ and KHK^-^/^-^ animals treated with vehicle (methylcellulose, purple) or a ketohexokinase inhibitor (KHKi, blue) (n= 6-8 mice per group). Data are presented as mean ± SEM; individual data points are overlaid. *P* values were calculated using one-way analysis of variance (ANOVA) with Dunnett’s multiple comparison test or two-way ANOVA with Tukey’s multiple comparison test.

**Supplementary Figure 2.**
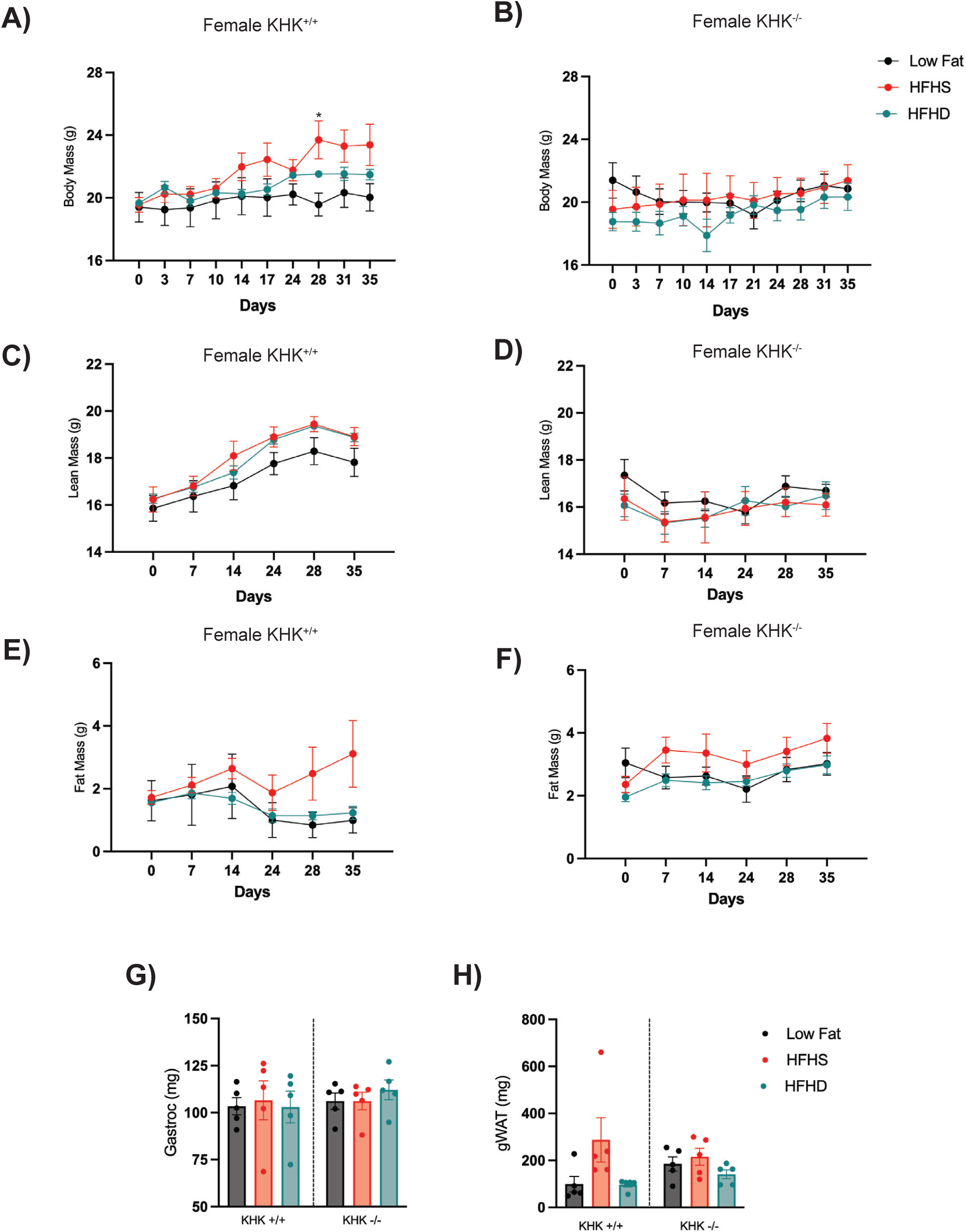
Dietary sucrose promotes weight gain in C57BL/6J female and can be prevented by KHK deletion. Female KHK^+/+^ and KHK^-/-^ animals received a low fat (black), high-fat high-sucrose (HFHS, red) or high-fat high-dextrose (HFHD, teal) diets for 5 weeks. **A)** Female KHK^+/+^ and **B)** Female KHK^-/-^ body mass over time. **C)** Female KHK^+/+^ and **D)** Female KHK^-/-^ lean mass over time. **E)** Female KHK^+/+^ and **F)** Female KHK^-/-^ fat mass overt time. After five weeks of different diet exposure, euthanasia was performed, and tissues such as **G)** Gonadal white adipose tissue (gWAT) and **H)** Gastrocnemius (gastroc) were collected and weighed. Data are presented as mean ± SEM; individual data points are overlaid. *P* values were calculated using two-way analysis of variance (ANOVA) with Tukey’s multiple comparison test or one-way with Dunnett’s multiple comparison test. * *p*=0.0336. *(Low Fat vs HFHS). n=5 mice per group.

**Supplementary Figure 3.**
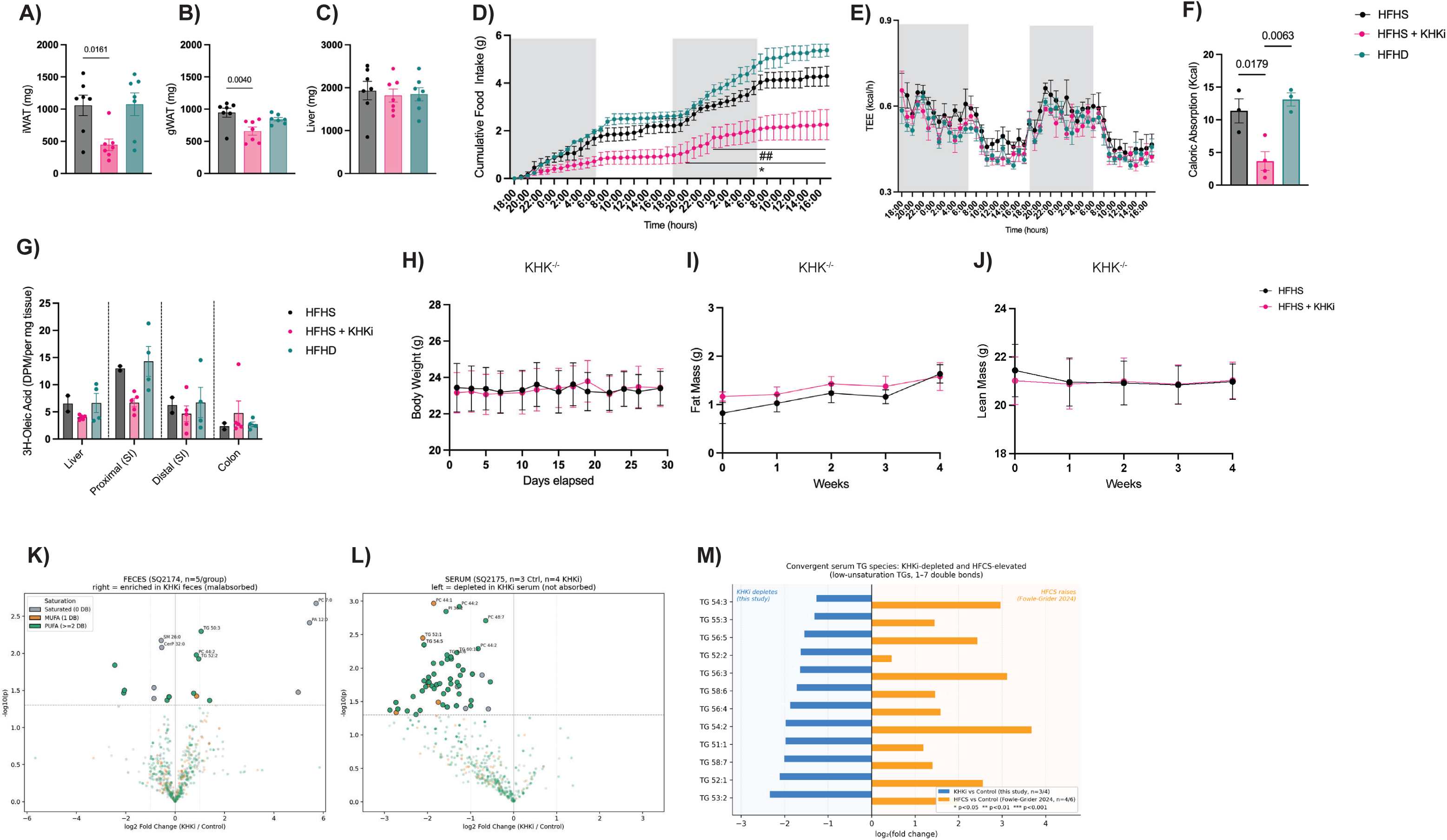
Small molecule PF-06835919 led to no changes in body weight in KHK^-/-^ mice under a HFHS diet: **A)** Inguinal white adipose tissue (iWAT), **B)** Gonadal white adipose tissue (gWAT) and **C)** Liver from obese mice after a month of oral treatment of PF-06835919 were collected at the end of the experiment (n=7 mice per group). **D)** Cumulative food intake (g) over a 48-hour period shows decreased energy consumption in KHKi group, when compared to HFHS (*) or to HFHD (^#^) group. **E)** Total energy expenditure (kcal/h) measured over 48 hours by indirect calorimetry reveals no significant differences among groups, with preserved circadian rhythmicity. **F)** KHKi treatment decreases caloric absorption (Kcal) compared to HFHS or HFHD groups (n=3 mice per group). **G)** ^3^H-Oleic acid levels were measured at the end of the experiment in different tissue sections such as liver, proximal intestine (small intestine = SI), distal intestine and colon in the different experimental groups (n=2-4 mice per group). No off-target effects were observed in **H)** Body weight, **I)** Fat mass and **J)** Lean mass over a period over a period of 4 weeks (n=8-9 mice per group) when KHK^-/-^ mice were treated with a HFHS diet + PF-06835919. HFHS: High-fat high-sucrose diet + vehicle, HFHD: high-fat high-dextrose diet + vehicle, HFHS + KHKi: High-fat high sucrose diet + PF-06835919. Volcano plots showing differentially abundant lipid species in **K)** feces (n=5 mice per group) and **L)** serum (n=3 control, n=4 HFHS + KHKi) of DIO mice fed a HFHS diet for 15 weeks. Each dot represents one lipid species, colored by degree of unsaturation. DB: double bonds, MUFA: monounsaturated, PUFA: polyunsaturated. The x-axis shows the log_2_ fold change (KHKi/Control) and the y-axis shows -log_10_ (*p*-value). **M)** Convergent serum TG species depleted by KHKi and elevated by HFCS (High fructose corn syrup). Mirror bar chart showing log_2_ fold changes for 12 triglyceride species (1-7 double bonds) that are significantly altered in opposite directions by KHK inhibition (blue, left; KHKi *vs*. control, n=3/4) and HFCS feeding (orange, right; HFCS *vs*. control, from Fowle-Grider *et al*^34^., n=4/6). Species are ordered by the magnitude of KHKi depletion. Data are presented as mean ± SEM; individual data points are overlaid. *P* values were calculated using one-way analysis of variance (ANOVA) with Dunnett’s multiple comparison test or two-way ANOVA with Tukey’s multiple comparison test. *: *p*< 0.05, **^/##^: *p<*0.001, ***: *p*<0.001.

**Supplementary Figure 4.**
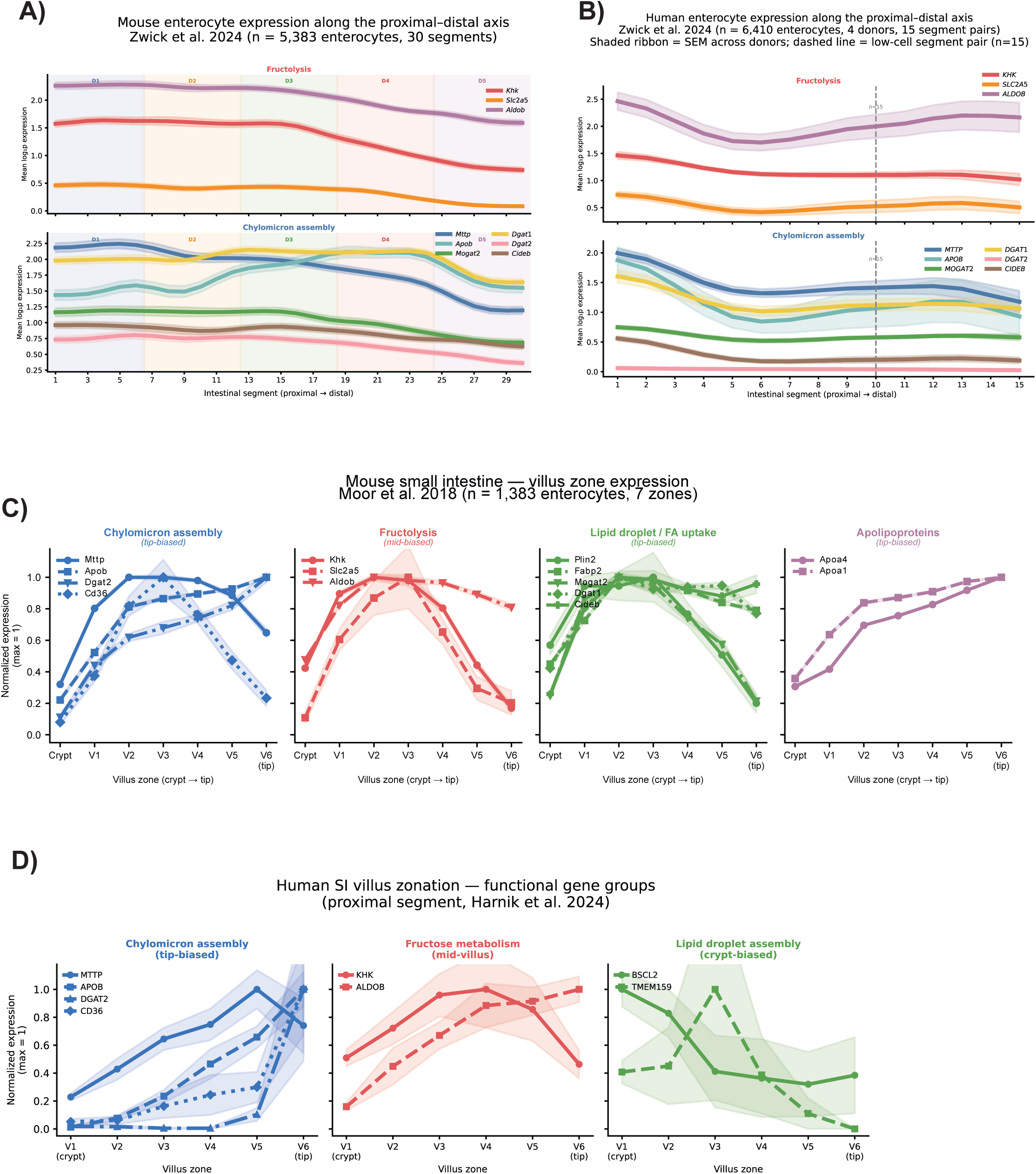
Villus zone-specific expression of functionally related fructolysis and lipid uptake/metabolism gene groups in mouse and human small intestine: **A)** Mouse and **B)** Human enterocyte expression of genes related to fructolysis and chylomicron assembly along the proximal-distal intestine axis. Value represents the mean scaled expression of each gene, colored by domain, with surrounding shade standard error bounds, across intestinal segments. Segment positions are numbered (x axis), and the positions of domain boundaries are noted with shade lines. Original data from Zwick *et al*.^9^ **C)** Normalized expression of genes involved in four lipid-handling functions across seven villus zones (crypt to tip) in mouse small intestinal enterocytes, replotted from Moor *et al.,*^35^. Chylomicron assembly genes (*Mttp*, *Apob*, *Dgat2*, *Cd36*; blue) and lipid droplet/fatty acid uptake genes (*Plin2, Fabp2, Mogat2, Dgat1, Cideb*; green) are enriched toward the villus tip. Fructolysis genes (*Khk, Slc2a5, Aldob*; red) peak at mid-villus zones (v2-v3). Apolipoproteins (*Apoa4, Apoa1;* purple) show a progressive tip-biased gradient. **D)** Normalized expression of orthologous human gene groups across six villus zones (crypt to tip) in the proximal small intestine, replotted from Harnik *et al*.,^8^ . Chylomicron assembly genes (*MTTP, APOB, DGAT2, CD36;* blue) are tip-based, fructose metabolism genes (*KHK, ALDOB;* red) peak at mid-villus zone (V3-V5) and lipid droplet assembly genes (*BSCL2, TMEM159;* green) are crypt-biased.

**Supplementary Figure 5.**
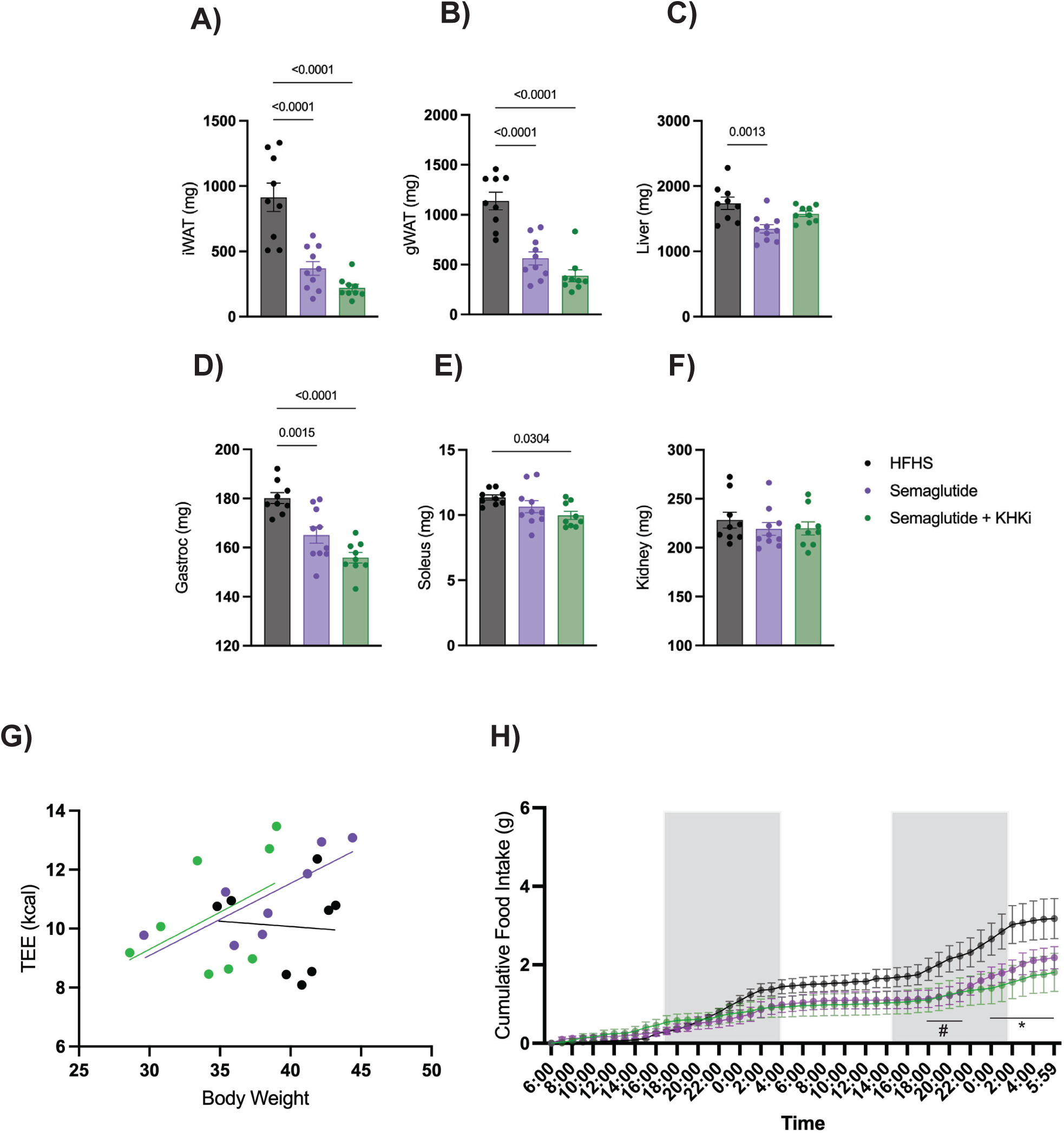
Semaglutide alone or in combination with PF-06835919 enhances fat and lean mass weight loss under a HFHS diet: At the end of the experiment, necropsy was performed and tissues such as **A)** iWAT (inguinal white adipose tissue), **B)** gWAT (gonadal white adipose tissue), **C)** Liver, **D)** Gastroc (Gastrocnemius), **E**) Soleus and **F)** Kidney were collected and weighted (n=9-10 mice per group)**. G)** Linear regression analyses of TEE (kcal) versus body weight (g) and **H)** Cumulative food intake (g) over the final 48 hours in metabolic cages (n=8 mice per group). Data are presented as mean ± SEM; individual data points are overlaid. *P* values were calculated using one-way analysis of variance (ANOVA) with Dunnett’s multiple comparison test. ^#/*^p=0.005, ^#^HFHS vs. semaglutide + KHKi, *HFHS vs. semaglutide

**Supplementary Figure 6.**
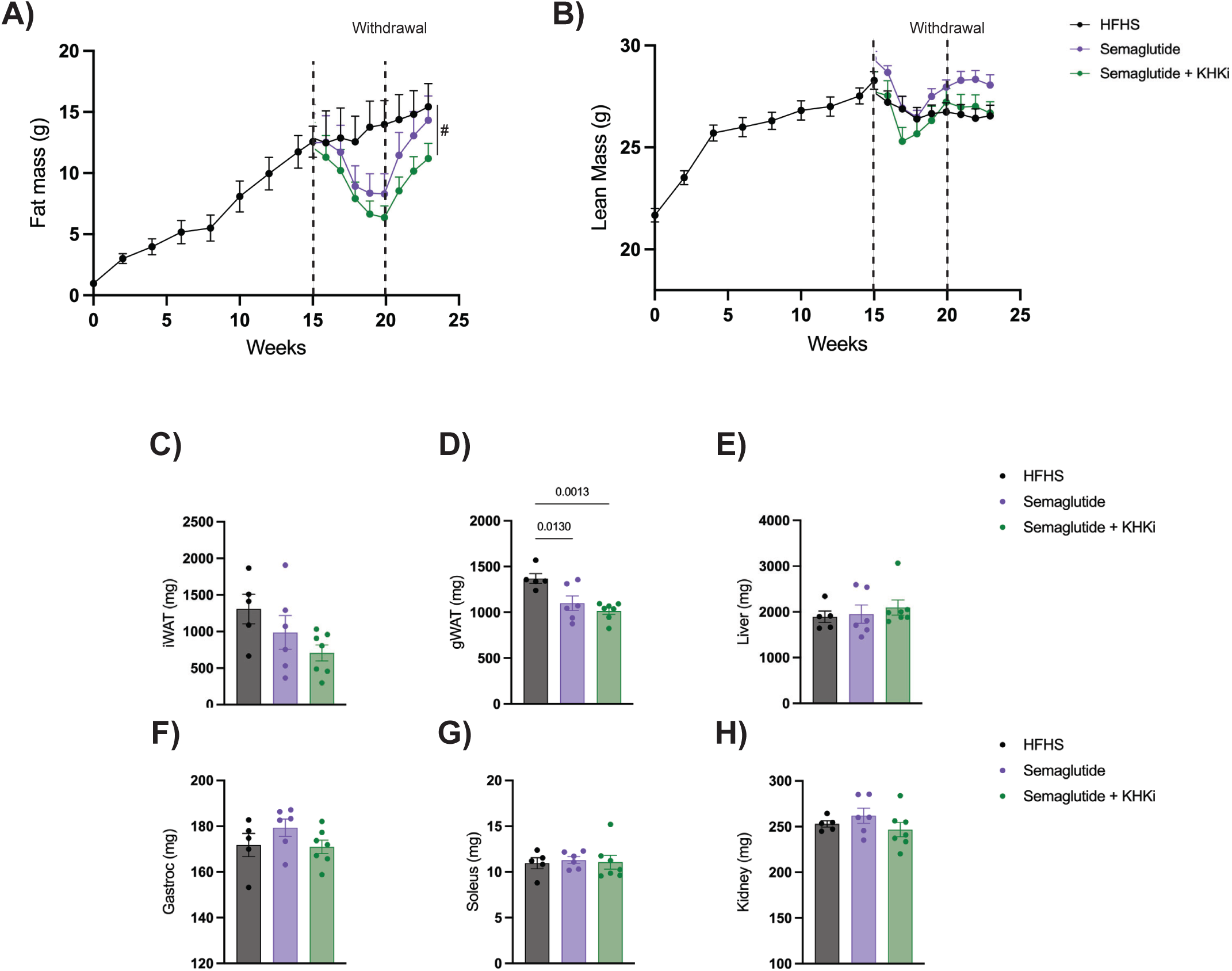
Small molecule PF-06835919 delays fat mass regains after semaglutide withdrawal: C57BL/6J male mice were fed a high-fat high-sucrose (HFHS; black) diet for 15 weeks to develop a DIO model. After this time, mice were randomized and subdivided in three different groups: continued HFHS (vehicle), HFHS + Semaglutide (30 nmol/kg; purple), or HFHS + Semaglutide + KHKi (PF-06835919; 50 mg/kg; green). **A)** Fat mass (g) and **B)** Lean mass (g) were measured longitudinally by body composition analysis. Tissue collection such as **C)** Inguinal white adipose tissue (iWAT), **D)** Gonadal white adipose tissue (gWAT), **E)** Liver, **F)** Gastrocnemius (gastroc), **G)** Soleus and **H)** Kidney from DIO mice after the withdrawal phase were collected at the end of the experiment (n=5-7 mice per group). Data are presented as mean ± SEM; individual data points are overlaid. *P* values were calculated using one-way analysis of variance (ANOVA) with Dunnett’s multiple comparison test. ^#^*p*=0.0215, ^#^HFHS vs Semaglutide+ KHKi.

**Supplementary Table 1:** Effects of HFHS feeding diet on female C57BL/6J and KHK^-/-^ mice.

|  | Female C57BL/6J |  |  | Female KHK<sup>-/-</sup> |  |  |
| --- | --- | --- | --- | --- | --- | --- |
|  | Low Fat | HFHS | HFHD | Low Fat | HFHS | HFHD |
| Insulin ng/mL | 0.98 ± 0.27 | 1.01 ± 0.25 | 1.19 ± 0.51 | 1.47 ± 0.50 | 0.49 ± 0.10 | 1.72 ± 0.59 |
| Leptin pg/mL | 1350.70 ± 485.64 | 5775.88 ± 2611.70 | 1686.46 ± 480.45 | 3064.44 ± 558.98 | 3942.42 ± 858.41 | 1870.06 ± 472.90 |
| Glucose (mg/dL) | 186.2 ± 7.86 | 252.2 ± 14.14** | 179.4 ± 5.89~~~ | 155.2 ± 8.40 | 134.2 ± 6.18 | 149.2 ± 11.27 |
| Triglycerides (mg/dL) | 46.28 ± 3.62 | 53.28 ± 4.36 | 48.07 ± 4.59 | 41.17 ± 6.28 | 48.76 ± 6.01 | 29.46 ± 2.35 |
| Cholesterol (mg/dL) | 56.43 ± 3.87 | 65.79 ± 9.30 | 79.63 ± 7.70 | 48.22 ± 2.19 | 57.56 ± 1.54 | 50.32 ± 7.13 |
| NEFA (mEQ/L) | 0.41 ± 0.06 | 0.52 ± 0.08 | 0.44 ± 0.06 | 0.30 ± 0.06 | 0.46 ± 0.08 | 0.38 ± 0.09 |

\*\*0.0014, ~~~0.0006. \*Low Fat vs HFHS, ~HFHS vs HFHD. Ordinary one-way ANOVA with Tukey's multiple comparisons test. Data are mean ± SEM. n= 4-5 mice per group

## Methods

### Animals and Diets

All animal studies were in accordance with NIH guidelines and approved by the Institutional Animal Care and Use Committee (IACUC) of the New York University (PROTO202400023). C57BL/6J male and female purchased from Jackson Laboratory were delivered at 8 weeks of age. Male and female *Khk^-/-^*mice lacking both KHK-A and KHK-C on the C57BL/6J background were provided by D. Bonthron (University of Leeds) and M. Lanaspa and R. Johnson (University of Colorado)^66^. The mice were housed in a specific pathogen-free facility with a strict 12:12 h light: dark cycle (06:00 am-18:00 pm), humidity at 45 ± 5% temperature of 22-24 °C and ad libitum access to food and water. Mice were regularly monitored for lethargy, gross weight loss, pallor and rectal prolapse.

For the weight gain trial, mice were randomized and fed with either a high-fat/high-sucrose diet (HFHS; 45% kcal from fat, 30% kcal from carbohydrate, 25% kcal from protein) (D19090602, Research Diets), a high-fat/high-dextrose diet (HFHD; 45% kcal from fat, 30% kcal from carbohydrate, 25% kcal from protein) (D19090601, Research Diets), or a low-fat diet (LF; 13% kcal from fat, 62% kcal from carbohydrate, 25% kcal from protein) (D17011901, Research diets) for 5 weeks. For the weight loss cohorts, mice were fed with HFHS diets for 15 weeks to induce obesity. Thereafter, mice were randomly allocated into five groups: control group (HFHS), KHKi group (HFHS + PF-06835919), semaglutide group (HFHS+ semaglutide), semaglutide + KHKi group (HFHS + semaglutide + PF-06835919) or HFHD group. Mice were oral gavage daily for 30 days with PF-06835919 (LabNetwork Inc) dissolved in 25% methyl cellulose (C9481, Sigma) at a concentration of 50 mg/kg or vehicle^31,32,67^. In the case of the GLP-1 therapy, semaglutide (HY-114118, MedChemExpress) was injected subcutaneously at a concentration of 30 nmol/kg every 3 days, respectively^63^.

Tracer studies were performed using custom Research Diets formulations (4.51 kcal/g, 45 kcal% fat, 30 kcal % carbohydrate, 25 kcal% protein). The carbohydrate fraction consisted of mixed sugars, with replacement of part of the carbohydrate content by isotopically labeled sugar. ^13^C-glucose diet (D25030702, Research Diets) contained 3.6 g/kg U-^13^C glucose and 13C-fructose diet (D25030703, Research Diets) contained 3.6 g/kg U-^13^C fructose. Both diets were otherwise matched to the high-fat/high-sugar control (D25030701, Research Diets).

### Measurement of glucose in blood

Glucose was measured using blood glucose meter (Accu-Chek Performa, Roche) from tail vein nick before necropsy in all the mice experiments.

### Serum Analysis

Blood was collected through cardiac puncture and stored on ice before centrifugation at 15,000 g for 10 min. Serum was subsequently flash-frozen and stored at -80°C before analysis. Small-scale, linear enzyme colorimetric assays were employed for measurement in duplicate of β-hydroxybutyrate (2440-58), triglycerides (2100-430), cholesterol (1010430) and nonesterified fatty acids (NEFAs) (999-34691) using the corresponding kit purchased from Fujifilm. Insulin levels were determined in duplicate by ultrasensitive mouse specific enzyme-linked immunosorbent assay (Crystal Chem, 90080) with intra-assay coefficient of variation (CV) <10%. Serum leptin levels (Alpco, 22-LEPMS-E01) and FGF21 levels (R&D, MF2100) were assayed in duplicates.

### Comprehensive metabolic monitoring

Mice were housed individually with ad libitum access to food and water 3 days prior to metabolic assessment to acclimate to housing conditions. All mice were acclimated to the metabolic cages for 24 h and then underwent metabolic monitoring for 48 h by indirect calorimetry using a Promethion high-definition multiplexed respirometry apparatus (Sable Systems, USA). Food intake and body mass were recorded continuously by gravimetric measurements within the cages. Energy expenditure was calculated using the Weir equation^68^. Energy expenditure is displayed as the total kcal per specified periods of time. For Whole-body fat oxidation rates were calculated from oxygen consumption (VO₂) and carbon dioxide production (VCO₂) obtained by indirect calorimetry. VO₂ and VCO₂ were expressed as mL·h⁻¹. Substrate oxidation was calculated using Frayn stoichiometric equations under the non-protein assumption. Fat oxidation (kcal·h⁻¹) as: FAT (kcal/h) = 0.00975 × (1.695 × VO₂ − 1.701 × VCO₂). Energy equivalents of 9.75 kcal·g⁻¹ for fat was applied. These equations derive from substrate oxidation stoichiometry and partition total gas exchange into fat contributions, assuming negligible protein oxidation.

### Stable isotope breath analysis in metabolic chambers

Mice were individually housed in Promethion metabolic phenotyping chambers (Sable Systems, USA), which continuously measure oxygen consumption (VO_2_), carbon dioxide production (VCO_2_), food intake and locomotor activity. To establish a natural-abundance isotopic baseline, animals were maintained on an unlabeled ^12^C control diet for 48 h before tracer exposure. On the experimental day, KHK^+^/^+^ and KHK^-^/^-^ mice received either vehicle (methylcellulose) or PF-06835919 (50 mg/kg, KHKi) by oral gavage approximately 2h before the onset of the dark cycle, generating four experimental groups: KHK^+^/^+^ + vehicle, KHK^-^/^-^ + vehicle, KHK^+^/^+^ + KHKi and KHK^-^/^-^ + KHKi. At the same time, the corresponding ^13^C- glucose or ^13^C-fructose tracer diet was placed into the chamber food hoppers. Food access remained blocked during this pre-loading period and was opened 1h before the lights off to synchronize tracer consumption with the beginning of the active feeding phase. Expired air from each chamber was continuously sampled by the integrated isotope gas analyzer while simultaneous VCO_2_ values were recorded by the Promethion system.

### Calculation of tracer oxidation rate

Expired ^13^CO_2_ enrichment was baseline-corrected using the mean signal from the final hour of the preceding unlabeled ^12^C diet period. Baseline-corrected ^13^CO_2_ enrichment was then combined with simultaneously measured VCO_2_ to calculate tracer-derived CO_2_ production and expressed as tracer oxidation rate (nmol/min). Data were aligned to the time of food access/gate opening.

### Normalization of oxidation to tracer intake

Food intake was continuously recorded from hopper mass measurements. Tracer intake was calculated from cumulative food consumption and the tracer concentration in the diet (4.0 g/kg). To account for inter-animal differences in feeding behavior and tracer exposure, tracer oxidation efficiency was calculated by normalizing cumulative tracer oxidation to tracer intake during a defined post-feeding analysis window. Oxidation efficiency was expressed as nmol tracer oxidized per mg tracer consumed.

### Body composition

Mice were weighed and body composition (Fat mass, lean mass, free water and total water) was measured using magnetic resonance spectroscopy using an EchoMRI-100H 2n1 with a horizontal probe (EchoMIR, Houston, TX).

### Fecal Bomb calorimetry

The feces from each mouse were collected over a 24 h period, during which mice were single-caged and housed at 22°C. Fecal pellets were dried at 60°C for 48h. The energy content of dry fecal samples was determinate with an adiabatic Parr 6200 Isoperibol Calorimeter equipped with a Parr 1109A Semimicro Oxygen Combustion Vessel (Parr Instrument Company, Moline, IL, USA) and calories were normalized to fecal weight as previously^69^.

### Fecal triglyceride content

Dried fecal pellets were weighted and fresh 30% alcoholic KOH was added to each sample (1:6). Feces were incubated at 60°C up to 6 hours for digestion. After digestion, the volume of the digested sample was measured and mixed with 1M MgCl_2_ (1:1.08), mixed and incubated on ice for 10 min. Centrifugation was carry for 30 min at 14000 rpm. Supernatant was used for triglyceride content described above.

### Serum and fecal lipidomics

Blood was collected through cardiac puncture from DIO mice 1 hour after an oral gavage of PF-06835919 or vehicle (methycellulose) and stored on ice before centrifugation at 15,000 g for 10 min. Serum was subsequently flash-frozen and stored at -80°C before lipidomics analysis. Fecal pellets were collected ad libitum from the last 3 days before the end of the experiment and stored at -80°C before lipidomics analysis. Serum samples were homogenized in a chloroform/methanol (2:1, *v/v*) and fecal samples were homogenized in an isopropanol/acetonitrile/water (4:3:1, *v/v*). Briefly, for serum samples, instrument mass accuracy was within tolerance (-3.0 ppm), LC column performance was stable (0.08 min RT range) and internal standard response variability was 55% across the samples and 19% across the internal standards alone, respectively indicating that sample extraction was minimally variable. For the fecal samples, instrument mass accuracy was within tolerance (-0.7 ppm), LC column performance was stable (0.4 min RT range) and internal standard response variability was 31% across the samples and 7% across the internal standards alone.

### Postprandial fat test

Mice were fasted for 12 h before receiving an oral gavage of PF-06835919 or vehicle (methycellulose). After an hour, mice received an oral bolus of 200 ul of lipid emulsion containing triolein, traced with [^3^H] triolein (MBq kg^-1^ 3 µCi) (PerkinElmer) or oleic acid, traced with [^3^H] oleic acid (MBq kg^-1^ 0.6 µCi) (PerkinElmer). After 0, 60, 120, 240, and 360 minutes, blood from the tail was extracted. Small intestine sections (proximal, distal and colon) were collected, weighed and dissolved in liquid scintillation cocktail (National Diagnostics, NC9186260), and radioactivity (in C.P.M) was measured by scintillation counting using a LS5000 scintillation counter (Beckman Counter).

### Small Histology and analysis

The intestine from each mouse was lateralized, washed with PBS-1X, opened, and swiss rolled. Small intestine tissue of mice was preserved in 4% paraformaldehyde solution in PBS-1X for 24 hours, followed by storage in 70% ethanol and embedded in paraffin. Five-micrometer sections were cut for hematoxylin and eosin (H&E) staining according to standard protocols followed by microscopy examination. Scanned H&E images of small intestine from each trial were downloaded as ScanScope Virtual Slide (SVS) files and divided into proximal small intestine, distal small intestine and colon. For villus length measurements, semi-automated analysis was performed. SVS files were opened in Image J and the length of the intestinal section was measured using the freehand measurement tool. Images were then stain-normalized to a standard H&E image using custom MATLAB (2024b) script using a method described previously^68^. A random image from the set was then loaded into the MATLAB Colour Thresholder and values were manually selected within the hue-saturation-value (HSV) colour space such that the intestinal villi, but not the other tissues such as lymph nodes or pancreas, were selected. These values were entered into a batch processing script that performed this villi segmentation on every image in the set. This resulted in binary images of pixels identified as villi and pixels identified as non-villi. The pixels occupied by villi were converted to area in um^2^ using the embedded scale from the original SVS file. The villi area was divided by the bowel segment length to yield the average thickness of the intestinal villi layer.

### Fluorescent *in situ* hybridization

Small intestine sections were mounted on SuperFrost Plus slides, air-dried at room temperature, and subsequently baked at 60°C overnight. For the simultaneous detection of various transcripts, we used fluorescent RNAscope (ACD; Advanced Cell Diagnostics Inc). All reagents were obtained by ACD unless specified otherwise. The pre-treatment protocol followed the manufacturer’s instructions and included an 8 min target retrieval at 98°C and 15 min incubation with Protease Plus. The detection protocol was performed with the RNAscope Multiplex Fluorescent v2 kit, following the manufacturer’s instructions. For each slide, RNAscope probes for bacterial dapB were used as negative control (320871, 1: 2000). The following catalog numbers were used for the detection of *Mttp* mRNA (899231,1:2000), and *Olfm4* mRNA, (311831,1:2000).

### Imaging and quantification of *in situ* hybridization

RNAscope sections were obtained using 20X objective (Ventra). Tile scans and Z-stacks were captured to cover the small intestine. To obtain quantification of *Mttp* mRNA expression from the ISH analyses along the crypt-villus axis and across proximal to distal intestinal domains, images were imported to Image J. Images were duplicated for proximal and distal intestine to create 8-bit images. Thresholds were set across all images. In brief, segmented lines were manually drawn on 10 villi in each domain of the intestine (proximal, middle, distal) per mouse and probe intensity was measured along this line from proximal to distal. Intensity was measured from the *Olfm4*+ crypt cells to the distal tip of the villi. In Rstudio (v.4.3.2), distance from crypt was normalized across villi and representative averages were generated.

### Electron Microscopy (EM)

Electron microscopy analysis of intestine samples was performed at the Microscopy Laboratory at NYU Langone Health. Fixation and preparation of samples for imaging by EM were completed as described previously^70^. Briefly, three C57BL/6J male mice treated with water or HFCS with DMSO or PF-06835919 were fasted overnight and then administered 200 uL of olive by oral gavage. After 120 min later, small intestines were collected. The small intestine was divided into domains A, C and the boundary between D/E^9^ (duodenum, jejunum, and ileum), and samples from each section were isolated and used for further processing. Samples were stained with 2% osmium tetroxide, dehydrated with ethanol, and embedded in Embed812 resin. Sem-thin section (1 mm) was cut and stained with 1% toluidine blue to evaluate the orientation of the sample. Ultrathin section (70 mm) was cut via ultramicrotomy, mounted on 200 mesh copper grids, and stained with lead citrate and uranyl acetate. Stained grids were imaged a STEM detector on a Zeiss Gemini300 Scanning Electron Microscope for low-magnification, whole-section imaging. Area of interested were randomly selected and imaged at higher resolution using JEOL1400 flash TEM microscope equipped with a Gatan Rio 16 camera (Gatan Inc. CA). All chemical and EM grids were purchased from Electron Microscopy Sciences (Hatfield, USA).

### TEM CLD analysis

Acquired TEM images were used to assess cytoplasmic lipid droplet (CLD) characteristics. Images from intact enterocytes representing the middle region of at least three villi per region per mouse were used for the analysis, resulting in the inclusion, on average, of ∼3 enterocytes per region per mouse for each treatment.

CLD diameter was measured using the Trainable Weka Segmentation^71^ v4.0 plugin running in Fiji. CLD area was estimated from the measured diameters using the formula area =π×(diam/2)^2^. Only CLD > 0.063 μm^2^ were considered. Electron density of CLDs were calculated following Cheng et al^38^. Electron microscopy images were analyzed using ImageJ, and the extracellular background as the internal standard for each picture. A total of 10 CLD from 10 different intact cells per animal were analyzed per condition.

### Reanalysis of published single-cell and spatial transcriptomic datasets

To characterize the spatial organization of fructose metabolism and lipid transport genes along the intestinal crypt-to-villus and proximal-to-distal axes, three published datasets were reanalyzed. Moor et al. 2018 (crypt-to-villus axis, mouse)^35^. Raw UMI count data and per-cell villus zone assignments from Moor et al. (2018) were obtained from Zenodo (DOI: 10.5281/zenodo.3403670). The dataset comprises 1,383 mouse jejunal enterocytes (27,998 genes) assigned to seven spatial zones (Crypt, V1–V6) using the authors’ landmark-gene spatial reconstruction algorithm. Raw counts were library-size normalized to counts per 10,000 (CP10K) and log1p-transformed (median library size: 6,754 UMIs per cell; range: 1,819–30,281). Zone cell counts were: Crypt, n = 235; V1, n = 247; V2, n = 168; V3, n = 68; V4, n = 312; V5, n = 209; V6, n = 144. Per-zone mean log1p(CP10K) expression and standard error of the mean (SEM) were computed for 14 target genes spanning fructolysis (Khk, Slc2a5, Aldob), chylomicron assembly (Mttp, Apob, Mogat2, Dgat1, Dgat2, Cideb), lipid droplet and fatty acid uptake (Plin2, Cd36, Fabp2), and apolipoproteins (Apoa4, Apoa1). For visualization, expression was max-normalized per gene (per-gene maximum = 1).

Zwick et al. 2024 (proximal-to-distal axis, mouse and human)^9^. Processed h5ad files from Zwick et al. (2024) were obtained from the CELLxGENE data portal (collection 3db5617e-2564-48ad-a981-d0b72a552248). The mouse dataset contains 19,847 cells × 16,966 genes from 30 consecutive proximal-to-distal small intestinal segments collected by MULTIseq barcoding; the human dataset contains 19,200 cells × 27,359 genes from 15 segment pairs across four donors (H1, H2, H4, H5). Expression values in the .X slot are log1p-normalized counts. Analyses were restricted to enterocytes: for mouse, cells annotated as ’Mature enterocyte’ (n = 3,311), ’Enterocyte’ (n = 1,072), or ’Enterocyte progenitor’ (n = 1,000) were retained (total n = 5,383; 95–339 cells per segment); for human, all enterocyte-annotated cells were retained (total n = 6,410; 15–1,056 cells per segment pair). Per-segment mean expression was computed for the same 14 target genes (mouse/human orthologues). For human data, per-donor per-segment means were computed and SEM was calculated across four donors. Smoothed expression curves were generated using a Gaussian filter (σ = 1.5 segments). Pearson and Spearman correlations between Khk/KHK and Mttp/MTTP expression profiles were computed across all segments.

Harnik et al. 2024 (crypt-to-villus axis, human)^8^. Pre-computed villus zonation tables from Harnik et al. (2024) were obtained from Zenodo (DOI: 10.5281/zenodo.11490477; file: S_table_zonation_by_segment.xlsx). The table provides per-zone mean expression and SEM for 20,023 genes across six villus zones (V1–V6, crypt to tip) in each of 15 intestinal segments from human donors, derived from spatially reconstructed single-cell RNA-sequencing data. Analyses were restricted to segment 1 (proximal intestine). Ten target genes were examined: chylomicron assembly factors (MTTP, APOB, DGAT2, CD36), fructose metabolism enzymes (KHK, ALDOB), and lipid droplet markers (BSCL2, TMEM159). Expression was max-normalized per gene for visualization. Center-of-mass (CoM) values, provided in the source table as the expression-weighted mean villus zone position (range 0–1, crypt to tip), were used to quantify spatial bias: KHK CoM = 0.50 (peak zone V4); ALDOB CoM = 0.64; MTTP CoM = 0.62; APOB CoM = 0.95; BSCL2 CoM = 0.36; TMEM159 CoM = 0.34.

All analyses were performed in Python 3.11 using pandas, numpy, scanpy, and matplotlib.

### Quantification and statistical analysis

All graphs show the mean and error bars represent the standard error of the mean (±SEM) as indicated in the figure legends. Statistical analyses were performed with Graphpad Prism 10 software, Python 3.11 or Rstudio (v.4.3.2). Sample normality was analyzed using one-way ANOVA or two-way ANOVA as indicated followed by Tukey’s multiple comparisons post hoc test. Significance was accepted when the *p-*value was lower than 0.05 (<0.05). Differences are marked with * or ^#^, indicating *^/#^*p*<0.05, **^/##^*p*<0.001, ***^/###^*p*<0.001 and ****^/####^*p*<0.0001 respectively, as specified in the figure legends. Each experiment was conducted with biological replicates and repeated no less than three times.

## Data availability

All data that support the findings of this study are available upon request from the corresponding author.

